# The genome of *Gallaecimonas pentaromativorans* strain 10A, isolated from a Pacific oyster, reveals the potential for hydrocarbon degradation and CRISPR-Cas defense

**DOI:** 10.1101/2024.10.01.616199

**Authors:** Yasmine Gouin, Adam Wilcockson, Kevin Xu Zhong, Amy M. Chan, Curtis A. Suttle

## Abstract

Bacteria in the genus *Gallaecimonas* are known for their ability to breakdown complex hydrocarbons, making them of particular ecological and biotechnological significance. However, few species have been isolated to date, and their ecological distribution has yet to be examined. Here, we report a novel bacterium in the genus *Gallaecimonas*, designated as *G. pentaromativorans* strain 10A, which was isolated from a Pacific oyster (*Magallana gigas*, a.k.a. *Crassostrea gigas*) collected from a farm experiencing a mass mortality event in British Columbia, Canada. *G. pentaromativorans* strain 10A is a rod-shaped, motile bacterium and has a circular genome of 4,322,156 bp encoding 3,928 protein-coding sequences (CDS). Phylogenetic analysis showed that strain 10A is closely related to members of *G. pentaromativorans.* Like other *Gallaecimonas* members, strain 10A harbours specific pathways involved in degrading polycyclic aromatic hydrocarbons (PAHs) and xenobiotic compounds, producing biosurfactants, and assimilating nitrate and sulfate; however, it is uniquely equipped with an additional 166 genes belonging to 147 protein families, including a putative *higB*-*higA* that likely contributes to enhanced stress response. Strain 10A also possesses clustered regularly interspaced short palindromic repeats (CRISPR) and CRISPR-associated (Cas) systems (CRISPR-Cas), prevalent in *Gallaecimonas* (detected in three out of four species), implying a potential defense mechanism against exogenous mobile genetic elements such as plasmids and viruses. We also mined publicly available databases to establish the widespread distribution of bacteria in the genus *Gallaecimonas* in seawater, sediments, and freshwater across latitude, suggesting its versatility and importance to environmental processes. Ultimately, this study demonstrates that the genome of *Gallaecimonas pentaromativorans* strain 10A, isolated from a diseased Pacific oyster, encodes for a suite of putative functions, including oil degradation, xenobiotic breakdown, biosurfactants production, assimilatory nitrate and sulfate reduction, and viral defense. This plasticity and breadth in metabolic function help to explain the cosmopolitan distribution of members of this genus.

## Introduction

Bacteria in the genus *Gallaecimonas* (class *Gammaproteobacteria*) are known for their ability to degrade polycyclic aromatic hydrocarbons (PAHs) and xenobiotic compounds, as well as produce biosurfactants [1–5]. Consequently, they have become important allies in bioremediation, particularly in the degradation of oil compounds and pollutants, including PAHs, toluene, fluorobenzoate, and chlorobenzene [1–5]. Furthermore, they also produce biosurfactants, natural surface-active molecules that facilitate processes such as emulsification, dispersion, and solubilization; thus, they are deployed in numerous industries, including agriculture, pharmaceuticals, and environmental remediation [2]. Additionally, one *Gallaecimonas* species has demonstrated antimicrobial activity against the bacterial pathogen *Vibrio harveyi* through the production of cyclic peptides, specifically diketopiperazines [6]. Despite their ecological and industrial importance, there are only four described isolates in the genus *Gallaecimonas*, each belonging to a four different species: *G. pentaromativorans* [1], *G. xiamenensis* [3], *G. mangrovi* [4], and *G. kandeliae* [5]. Members of the genus have been reported from mid-latitude intertidal sediments and seawater, as well as in symbiotic associations with plant hosts, including the mangrove *Kandelia obovate* [1, 3–5]. In spite of these discoveries, the number of cataloged *Gallaecimonas* species remains limited, and their ecological distribution is largely unexplored.

Here, we genomically characterize *Gallaecimonas pentaromativorans* strain 10A, which we isolated from a Pacific oyster during a mass mortality event of oysters in British Columbia, Canada. This study benchmarks the genomic architecture and genetic capabilities of strain 10A against those of other members within the genus *Gallaecimonas*. Additionally, by interrogating publicly accessible datasets of 16S ribosomal RNA (rRNA) gene sequences, we elucidate the global distribution of members of the genus *Gallaecimonas*. This research highlights the potential role of these bacteria as biosurfactant producers and oil-like pollutant degraders in oysters.

## Materials and Methods

### Bacterial isolation and culture conditions

Pacific oysters were collected from a tray at an aquaculture facility in the Baynes Sound area (49.5078° N, 124.8272709° W), British Columbia, Canada, during a mortality event in July 2020. These oysters exhibited varying degrees of mortality (non-gaping to gaping, dead); shell lengths ranged from 5 to 7 cm. The oysters were opened to harvest tissue samples within 1 hour of collection. Samples were placed into sterile Whirl-Pak sample bags and immediately flash frozen in a liquid nitrogen (LN) dry shipper, where they remained frozen in LN vapors for one week, then stored at -80 °C. *G. pentaromativorans* strain 10A was isolated from tissue collected from a non-gaping oyster. Approximately 1.5 grams of frozen oyster tissue was aseptically transferred to a sterile 50 mL Falcon tube, thawed on ice, and then combined with 1.5 mL of sterile F/2 seawater media [7]. This mixture was gently vortexed to create a homogenate and used to isolate bacteria.

To prepare spread plate cultures, 50 microliters of the oyster homogenate was pipetted onto CPM-24 plates (0.05% Difco Casamino Acids, 0.05% Difco Peptone, 1% Fisher Scientific purified agar; prepared with 24 practical salinity units (PSU) seawater) [8, 9] and MLB-24 plates (CPM- 24 with additional 0.05% Yeast Extract and 0.3% glycerol) [8, 9] and sterile 10-uL plastic inoculating loops are used to evenly spread the sample over the plate surface until all liquid is adsorbed. After incubating at room temperature (ca. 21°C) for about a week, a colony was cleanly picked and re-streaked four times onto CPM-24 plates, using a single well-separated colony each time in order to obtain axenic clonal cultures. Subsequently, the purified isolate was routinely cultured with MLB media prepared with 24 PSU seawater (MLB-24); stock cultures were preserved in 20% (v/v) glycerol and stored at -80 C.

The soft agar motility test [10] was performed by stabbing a tube of MLB-24 soft agar (0.6% agar) with cells picked from a colony grown on MLB-24 agar and monitoring growth away from the stab line after 2 to 3 d.

### Transmission electron microscopy (TEM)

To assess the morphology of strain 10A, a few colonies from a culture grown on MLB-24 agar were resuspended in 0.2 µm-filtered seawater, then fixed with 0.2 µm-filtered EM grade glutaraldehyde (25%) to achieve a final concentration of 1% glutaraldehyde. The fixed sample was adsorbed to the shiny side of a formvar-carbon 400 mesh copper grid (Cat# 01754-F, TedPella, CA) for 5 min. Excess sample was wicked away with filter paper, and the grid was then stained with 2% uranyl acetate for 30 s. Excess stain was wicked away, and the grid was allowed to air dry at room temperature for at least 5 minutes before visualization at 120 kV on a Tecnai Spirit transmission electron microscope.

### Genomic DNA extraction, sequencing, and genome assembly

To prepare a sample for whole genome sequencing, strain 10A cultures grown on MLB-24 agar plates for 2 d were used to inoculate two 50 mL Falcon tubes, each with 20 mL of MLB-24 broth. The cultures were grown in an orbital shaker at 24°C and 150 rpm for 2 d. Cells were harvested by pelleting in a Beckman Allegra X-22R centrifuge with a swinging bucket rotor at 3730 x *g* and 10°C for 15 min. The cell pellets were resuspended in approximately 4 ml of sterile F/2 seawater media (24 PSU) and dispensed into two 2 mL cryovials. Cells were pelleted in a Beckman Allegra X-22R centrifuge with a fixed angle rotor at 9000 x *g* and 10°C for 5 min, then frozen using dry ice after removal of the supernatant.

The bacteria were submitted to the Microbial Genome Sequencing Center (MiGS) at the University of Pittsburgh (Pittsburgh, PA) for genomic DNA extraction (Zymo fungal/bacterial DNA miniprep kit; Zymo Research, Irvine, CA), and hybrid assembly sequencing (Small Nanopore Combo package). The Illumina sample was prepared using the Illumina DNA library prep kit (Illumina, Inc., San Diego, CA) and sequenced on an Illumina NextSeq2000 instrument with 151-bp paired-end chemistry, generating 4,117,269 Illumina short-reads. The Nanopore sample was prepared using the Oxford Nanopore Technologies (ONT, UK) ligation sequencing kit and sequenced on a MinION instrument using an R9 flow cell (R9.4.1), with base calling performed using ONT Guppy v.4.2.2, yielding 144,908 ONT long-reads. Adapters and low-quality reads were trimmed using bcl2fastq v.2.19.0 [11] and porechop v.0.2 [12] for Illumina and ONT sequences, respectively. Hybrid assembly with Illumina and ONT reads was performed using Unicycler v.0.5 [13]. The integrity of the bacterial genome was checked using CheckM v.1.0.18 [14].

### Taxonomical classification and phylogenetic analysis

The taxonomic identity of strain 10A was determined using Kaiju v.1.8.2 [15], CAT v.5.2.3 [16], MetaErg v.1.2.3 [17], the NCBI Prokaryotic Genome Annotation Pipeline (PGAP) v.6.0 [18], and GTDBtk v.2.3.2 [19] with default parameters. Once identified as a member of the genus *Gallaecimonas*, phylogenetic placement was done by whole-genome phylogeny using GTDBtk v.2.3.2 [19] and publicly available genomes of *Gallaecimonas* spp. from the Genome Taxonomy Database (GTDB, v.217), using two strains of *Escherichia coli* (GTDBtk accession numbers: RS GCF 011881725.1 and RS GCF 003697165.2) as an outgroup. Phylogenetic analysis of the 16S rRNA gene was also done on complete 16S rRNA sequences for *Gallaecimonas* spp. from the SILVA rRNA gene database v.138.1 [20], and rooted with sequences from two strains of *Escherichia coli* (NCBI accession number: CP033092 and MT215717) as an outgroup. These sequences, along with that of strain 10A, were aligned using MAFFT v.7 [21] with default parameters. The alignments were then trimmed using Gblocks [22] with default parameters. The maximum-likelihood tree was constructed using PhyML 3.0 [23] with a bootstrap parameter of 100, and the best model (TN93) was determined using MEGA11 [24]. The tree was visualized with Geneious Prime v.2023.2.1.

### Gene prediction, annotation, and visualization

The genome was annotated using the NCBI Prokaryotic Genome Annotation Pipeline (PGAP) v.6.0 [18] with default parameters. Subsequently, the PGAP-predicted genes were annotated using eggNOG-mapper v.2 [25] with default parameters based on ortholog mapping against the eggNOG (Evolutionary genealogy of genes: Non-supervised Orthologous Groups) database, providing functional annotations such as seed ortholog (i.e., gene family), eggNOG OGs, COG category, PFAMs (i.e., protein family), EC, GO terms, CAZy, BiGG Reaction, BRITE, and KEGG profiles [26] including KEGG Ortholog (ko), KEGG Pathway, KEGG Module, KEGG Reaction, KEGG rclass, and KEGG TC. In addition, MetaErg v.1.2.3 [17] was also employed using default parameters to provide supplementary annotation, including MetaCyc pathways. Genome features were visualized using Proksee [27], and their KEGG profiles were visualized using FuncTree v.0.8.4 [28]. Additionally, CRISPR spacer sequences were identified using CRISPRFinder v.1.1.2 [29] with default settings.

### Comparative analysis of *Gallaecimonas* genomes

The genome of strain 10A was compared with those from the following genomes for *Gallaecimonas* spp. in NCBI: *G. xiamenensis* strain 3-C-1 [3, 30], *G. pentaromativorans* strain CEE_131 (Leibniz Institute DSMZ culture collection ID: DSM21945) [1], *G. mangrovi* strain HK- 28 [4], and *G. kandeliae* strain Q10 [5]. A pan-genome analysis and visualization were performed using PGAP v.6.0, eggNOG-mapper v.2, R packages ggplot2 v.3.4.4 [31] and ggvenn v.0.1.10 [32], and Proksee [25] with default settings. Prior to the analysis, to ensure consistent gene calling and annotation, the genomes were re-annotated using PGAP v.6.0 and eggNOG-mapper v.2 following the methodologies outlined above in “Gene prediction, annotation and visualization”. PGAP-predicted genes were grouped into different gene families (i.e., seed_ortholog) based on eggNOG-mapper annotations to determine shared genes across the different genomes. The core genomic features, including gene families (i.e., seed_ortholog), protein families (i.e., PFAMs), CAZy, COG profiles, EC, eggNOG Ogs, GO terms, KEGG ko profiles, and KEGG Module profiles, were defined by the presence of identical IDs across the analyzed genomes. Comparisons of genomic features (e.g., genes and their associated features such as gene families (i.e., seed_ortholog), protein families (i.e., PFAMs), CAZy, COG profiles, EC, eggNOG Ogs, GO terms, and KEGG profiles) were analyzed and visualized using R packages ggplot2 v.3.4.4 [31] and ggvenn v.0.1.10 [32]. Key functional pathways found in strain 10A were depicted using BioRender (https://www.biorender.com).

### Ecological distribution of *Gallaecimonas*

The ecological distribution (presence/absence and relative abundance) of *Gallaecimonas* spp. was investigated based on 16S rRNA gene sequences between 2003 and 2019 obtained from the Global Biodiversity Information Facility (GBIF) database [33] (S4 Table). Using the coordinates of the sampling sites provided in occurrence data, the global distribution was visualized using R packages ggplot2 v.3.4.4 [31], sf v.1.0-14 [34], and maps v.2.3-2 [35]. Additionally, the proportion of the number of 16S rRNA gene sequences assigned to *Gallaecimonas* spp. across different environments was determined through the habitat types of the sampling sites.

## Results

### Isolation of strain 10A

*Gallaecimonas pentaromativorans* strain 10A was isolated from an oyster aquaculture farm experiencing a mortality event. The cultures were purified by streaking repeatedly on solid media. When grown on MLB-24 plates, at 21°C for 3-4 days, colonies were circular, convex, colorless and translucent, and 0.5- to 1- mm in diameter. Colonies that are well separated from each other can be as large as 3 mm in diameter in 3-week-old cultures; these colonies appear to be less translucent and whiter in color. Strain 10A cells stained with 2% uranyl acetate and visualized with transmission electron microscopy (TEM) were rod-shaped, approximately 0.5-μm wide, and up to 2-μm long, and some appeared to possess a polar flagellum. A soft agar motility stab test was done by stabbing a tube of soft agar with cells. Diffuse waves and swirls of cell growth away from the initial stab line confirmed that strain 10A was motile.

### Genomic characterization of the strain 10A

Hybrid sequencing using Illumina and Nanopore technologies allowed for assembling the genome sequence of strain 10A, which consisted of a circular chromosome of 4,322,156 bp with an average GC-content of 58.3% (Fig 1). The integrity of the bacterial genome was validated using CheckM v.1.0.18, which showed that the genome was 99.82% complete and had 0.00% contamination compared to 899 reference gammaproteobacterial genomes in the database. Annotation using the National Centre for Biotechnology Information (NCBI) Prokaryotic Genome Annotation Pipeline (PGAP) v.6.0, revealed that strain 10A contains genes coding for 18 rRNA (six of each 5S, 16S, and 23S rRNAs), 86 tRNAs, and four ncRNAs, as well as 3,928 protein-coding sequences (CDS). These CDS belong to 2734 protein families (i.e., PFAMs) and encode proteins involved in a wide array of biological processes integral to metabolic and regulatory functions. Putatively, they include 21 Clusters of Orthologous Genes (COGs), 891 enzymes, 2,564 Gene Ontology profiles (GO), 1,359 KEGG Ortholog (ko) profiles, as well as 112 KEGG and 439 MetaCyc pathways (Fig 1; S1-2 Figs).

**Fig 1.**
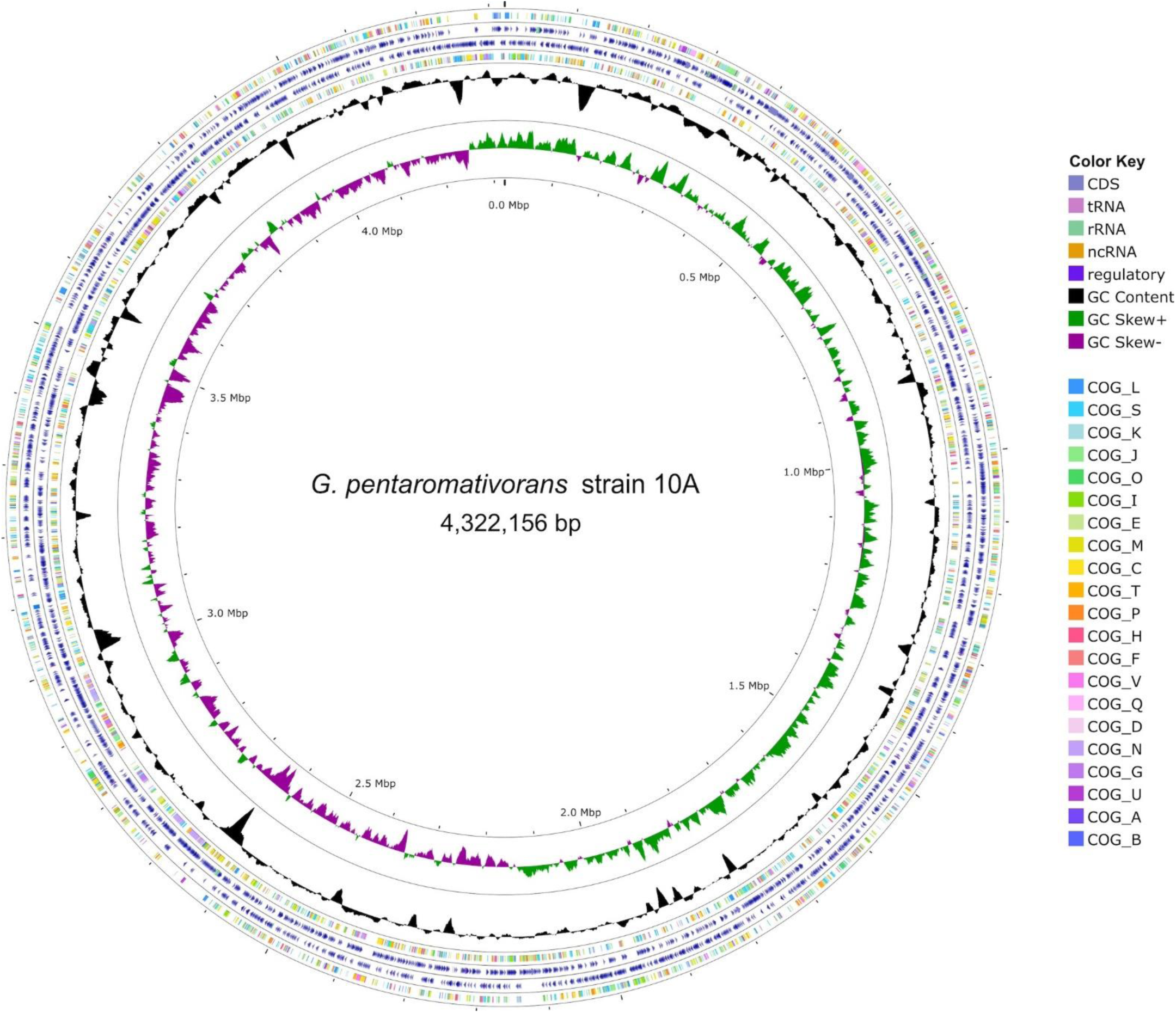
The complete genome of the *Gallaecimonas pentaromativorans* strain 10A. The outermost to innermost rings of the map represent the following: clusters of orthologous genes (COGs) functional categories for forward strand coding sequences; forward strand sequence features; reverse strand sequence features; COGs functional categories for reverse strand coding sequences; black ring shows GC content; and GC skew, with the green and purple bands representing positive and negative values, respectively.

Phylogenetic analysis of the 16S rRNA gene sequences available in the SILVA rRNA gene database v.138.1, showed that strain 10A clustered with members of *G. pentaromativorans* with a high bootstrap value (87/100; Fig 2A). Phylogenomic analysis based on whole genomes of *Gallaecimonas* spp. are consistent with strain 10A being closely related to *G. pentaromativorans* strain CEE_131 (Leibniz Institute DSMZ culture collection ID: DSM21945; Fig 2B). Indeed, between strain 10A and strain CEE_131 (DSM21945), the average nucleotide identities (ANI) of the genome and 16S rRNA gene are 98.98% and 97.3% similar, respectively (Fig 2), consistent with 10A and CEE_131 (DSM21945) being the same species. This classification is further supported based on overall genomic features, which place strain 10A in the genus *Gallaecimonas* (using NCBI PGAP v.6.0, GTDBtk v.2.3.2, MetaErg v.1.2.3 and CAT v.5.2.3) or in the species *G. pentaromativorans* (using Kaiju v.1.8.2).

**Fig 2.**
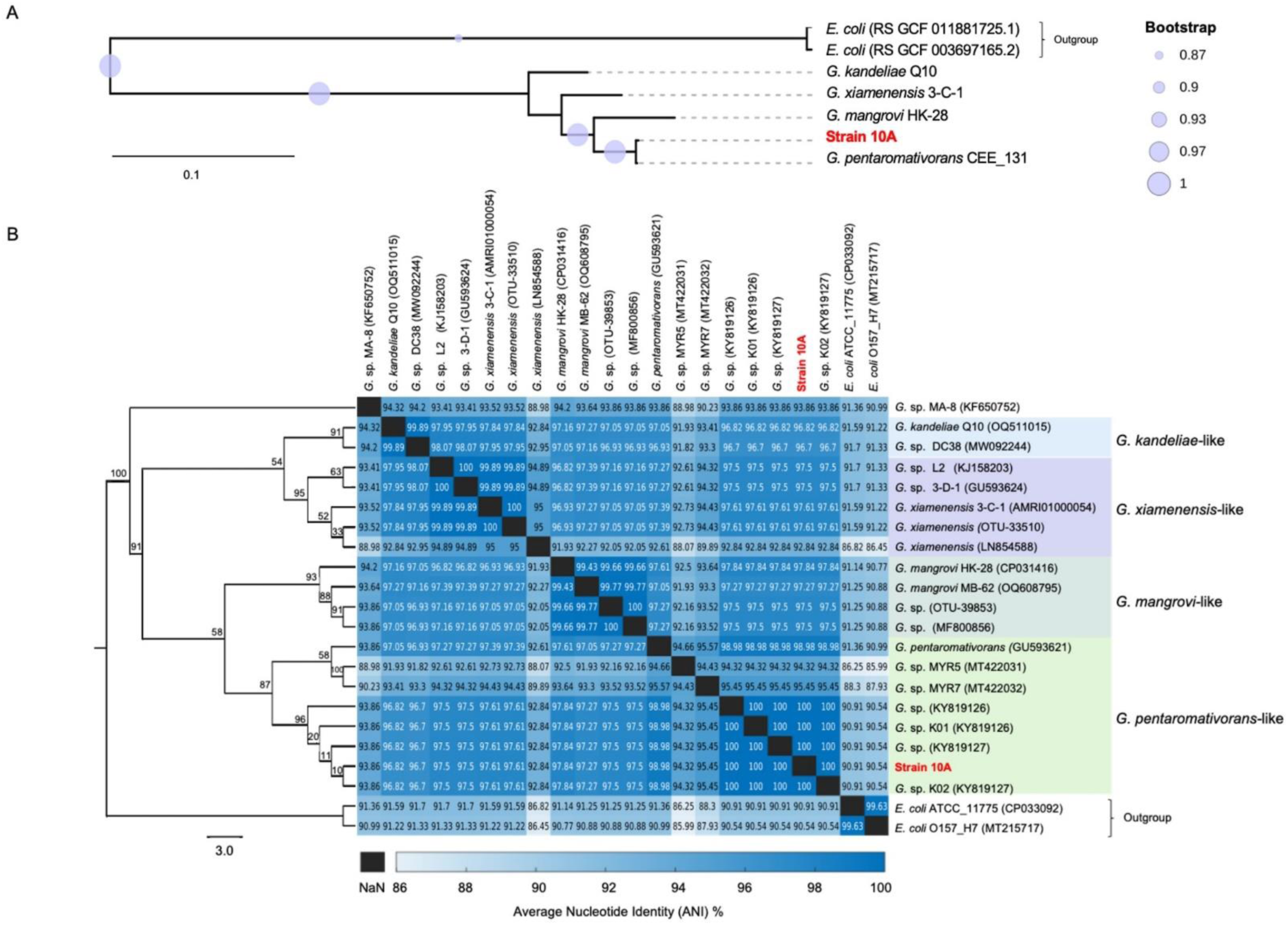
Phylogenetic relationship among DNA sequences from *Gallaecimonas* spp. (A) Phylogenomic tree of bacteria in the genus *Gallaecimonas* based on data in NCBI Reference Sequence Database (RefSeq, v.216). The tree was built based on the overall genome sequences and rooted with two genome sequences from *Escherichia coli* (GTDBtk accession numbers: RS GCF 011881725.1 and RS GCF 003697165.2) as an outgroup. (B) Phylogenetic relationship of the 16S rRNA gene sequences from strain 10A and other *Gallaecimonas* spp. found in the SILVA rRNA gene database v.138.1. The maximum-likelihood tree was built using 100 replicates and rooted with sequences from two strains of *Escherichia coli* (NCBI accession number: CP033092 and MT215717) as an outgroup. The value in the heatmaps associated with the phylogenic tree represents the average nucleotide identity (ANI) of 16S rRNA gene sequences between two strains on each of the x and y axes.

### Comparative analysis of *Gallaecimonas* genomes

The shared genomic features among isolates of *Gallaecimonas* spp. were investigated by a pan- genomic analysis of *G. xiamenensis* strain 3-C-1, *G. pentaromativorans* strain CEE_131 (DSM21945), *G. mangrovi* strain HK-28, *G. kandeliae* strain Q10, and *G. pentaromativorans* strain 10A (Fig 3A-B). These five isolates revealed 5562 genes; 38.1% were designated as “core” genes because they are shared by all the isolates, while the remainder are distributed across species. The “core” genes, which represent 60.45 to 66.95% of the genes in each species, were integrated into 1677 gene families (i.e., seed_ortholog in eggNOG-mapper annotation), 1586 KEGG ko profiles, 230 KEGG Modules, and 401 KEGG Pathways, as predicted using eggNOG-mapper (Fig 3B; S1-2 Figs; S2 Table). These “core” genes are involved in metabolic processes such as cell growth and death, binary fission, the citrate cycle, the metabolism of amino acids (e.g., *HisD, IlvC, SerA, Tdh),* and peptidoglycan synthesis (i.e., *murABCDEF, Pbp* genes*, Ddl)*. Notably, several highly conserved pathways, with over 50% coverage for both KEGG module and pathway profiles, entail two-component systems crucial for environmental processing: osmotic stress response (*MtrB-MtrA)*, cell fate control (*PleC-PleD)*, type 4 fimbriae synthesis *(PilS-PilR)*, cPHB biosynthesis (*AtoS-AtoC)*, membrane lipid fluidity *(DesK-DesR)*, capsule synthesis *(ResC-ResD- ResB)*, hexose phosphate uptake (*UhpB-UhpA)*, and cell wall metabolism *(VicK-VicR)* (S1 and S2 Figs).

**Fig 3.**
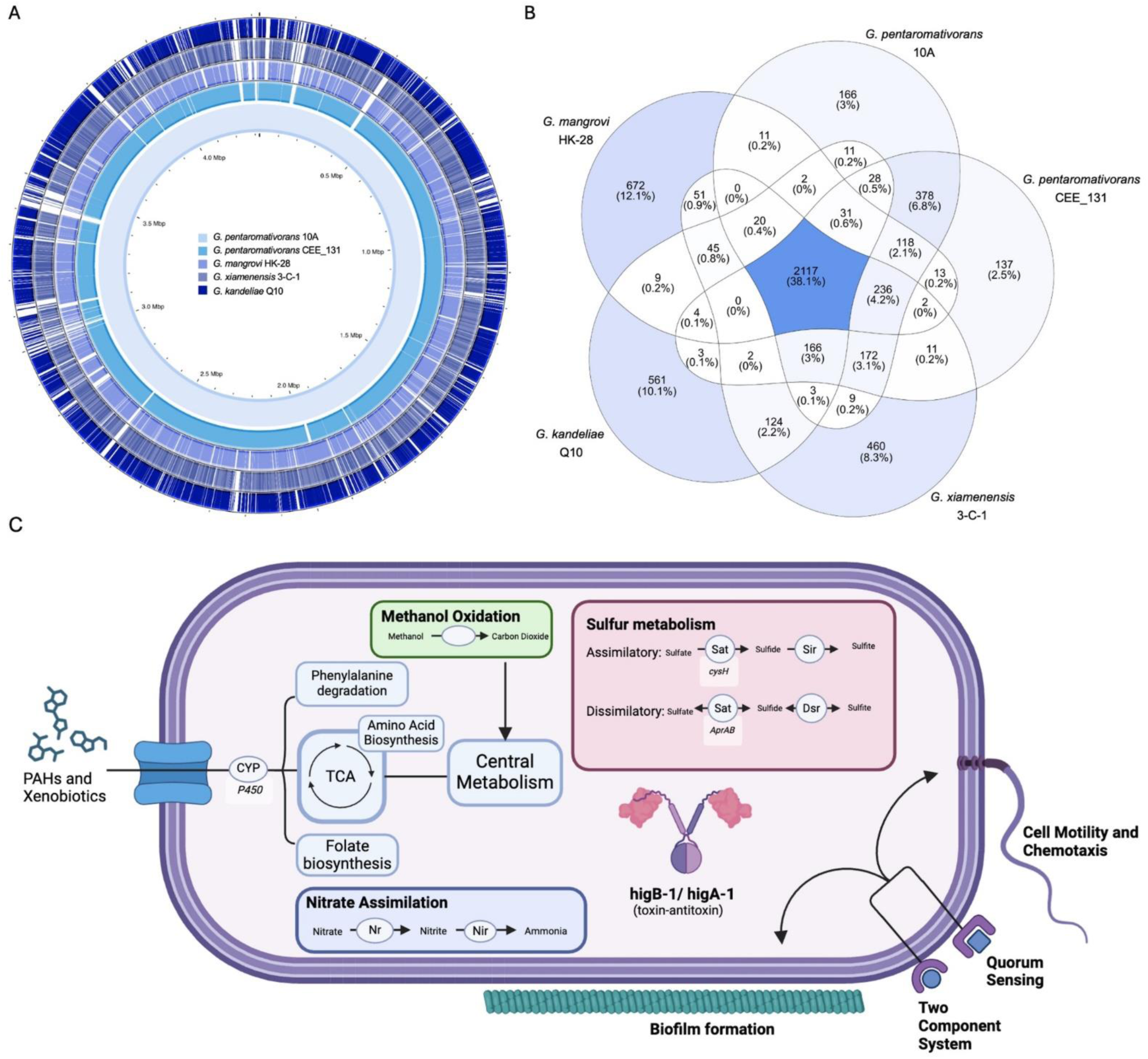
Comparative genomics among strains of *Gallaecimonas* spp. (A) Comparative map depicting genomic arrangements of four other isolates of *Gallaecimonas* spp. to strain 10A. (B) Venn diagram illustrating the distribution of shared and unique genes among the five strains of *Gallaecimonas* spp. (C) Key functional pathways found in *G. pentaromativorans* strain 10A, including PAHs and xenobiotic degradation, methanol oxidation, nitrate assimilation, sulfur metabolism, higB-1/higA-1 toxin-antitoxin system, and biosurfactant-producing pathways.

Despite the conserved genetic nature among isolates of *Gallaecimonas* spp., each member has between 137 and 672 characteristic genes, exhibiting inter-species genomic variability (Fig 3B; S2 Table). For instance, *G. xiamenensis* strain 3-C-1 bears 460 distinct genes (13.5% of genes in strain 3-C-1), among which are those related to copper tolerance, such as multicopper oxidase, *CusS-CusR* two-component system and divalent heavy-metal cations transporters (*Zip*). In contrast, *G. kandeliae* strain Q10 possesses 561 specific genes (18.1% of its genes), including those encoding zinc-binding dehydrogenase (ADH_N, ADH_zinc_N, ADH_zinc_N_2) and AMP-dependent synthetase and ligase (*AMP-binding, AMP-binding_C*). Similarly, *G. mangrovi* strain HK-28 carries 672 different genes (17.7% of its genes), encompassing those involved in the ABC transporter superfamily (*ABC_tran, oligo_HPY*), and guanosine tetraphosphate metabolism (*RelA, SpoT*). Even *G. pentaromativorans* strains CEE_131 (DSM21945) exhibits 137 specific genes (4% of its genes), including cysteine-rich domain-containing proteins (CCG) and Type I restriction enzyme R protein N terminus (*HSDR_N, HSDR_N_2*).

Intraspecies conservation of genes is evident in *G. pentaromativorans*, represented by strains CEE_131 (DSM21945) and 10A. These isolates share 378 genes, representing 40 KEGG ko profiles, that differ from those found in other *Gallaecimonas* spp. (S2 Fig; S2 Table). Of note are genes encoding aldehyde dehydrogenase (e.g., *badH)*, thiamine pyrophosphate enzymes (e.g., *poxB)*, alpha/beta hydrolases, SMART protein phosphatase 2C domain proteins, and fatty acid desaturase (S2 Table).

Inter- and intra-species variation among isolates of *Gallaecimonas* spp. is evident through analysis of strain 10A. In comparison to other isolates of *Gallaecimonas* spp., including *G. pentaromativorans* strains CEE_131 (DSM21945), strain 10A uniquely possesses 166 genes (4.2% of genes in strain 10A), integral to 147 protein families, 14 ko profiles and 1 KEGG pathway (Fig 3B; S2 Fig; S2 Table). These genes are predicted to encode for proteins involved in diverse biological functions, including DNA methylation (K00590), heme export for c-type cytochrome biogenesis (*CcmD*), flagellar transcriptional activation (*FlhC*), peptidase activity (K06992), capsule biosynthesis (*hipA*) and hemolysin activity (*shlB).* Moreover, a putative higB-1/higA-1 toxin/antitoxin (TA) system (e.g., K21498) was found in strain 10A, but was absent in other *Gallaecimonas* strains (Fig 3B-C). In addition, a KEGG pathway consisting of (i) a two-component system (ko02020), (ii) quorum sensing (ko02024), (iii) biofilm formation (ko02026), and (iv) flagellar assembly (ko02040), was identified in strain 10A, but not in other *Gallaecimonas* strains (Fig 3B-C). While not unique to strain 10A, of particular note are vital marker genes involved in methanol oxidation (e.g., *pqq, xoxF, mxat*), as well as complete pathways for assimilatory and dissimilatory sulfate reduction (M00176, M00596) and nitrate assimilation (e.g., *nasA*). Further, analysis using eggNOG-mapper and FuncTree revealed that strain 10A exhibits the genetic capacity for xenobiotic and PAHs degradation and metabolism through enzymes, such as cytochrome P450, which targets a diverse range of compounds, including benzoate, ethylbenzene, caprolactam, chloroalkane and chloroalkene, atrazine, styrene, naphthalene, nicotinate, and nicotinamide (Fig 3; S3 Fig).

CRISPR-Cas systems, a well-known prokaryotic defense system against exogenous plasmids and viruses [36], were also discovered in members of *Gallaecimonas* (Fig 4; S4-8 Figs). *Gallaecimonas mangrovi* strain HK-28 and *G. pentaromativorans* strain CEE_131 (DSM21945) had identical *Cas*-TypeIF clusters, which included *cas3-cas2, csy1, csy3, cas1, csy2,* and *cas6* (Fig 4; S4-6 Figs). Furthermore, four Cas clusters were identified in *G. xiamenensis* strain 3-C-1 (Fig 4; S7 Fig). Of note, two distinct Type I Cas clusters, featuring cas3a and cas3, respectively, and a Cas-TypeIIIU cluster containing *Csx3*-TypeIIIU were detected. In addition, a TypeIE CRISPR-Cas system was also found in both *G. xiamenensis* strain 3-C-1 and strain 10A and was characterized by a *cas2, cas1, cas6, cas5, cas7, cse2, cse1*, and an accessory c*as3-*TypeI (Fig 4; S7-8 Figs). Moreover, three CRISPR arrays (i.e., including the CRISPR repeat and spacer sequences), two with one spacer, and one with 95 spacers, were detected in the genome of strain 10A (S8 Fig; S3 Table). In summary, between one and ten CRISPR arrays were detected in all five *Gallaecimonas* genomes (Fig 4; S4-S8 Figs), although no *Cas* genes were identified in *G. kandeliae* strain Q10.

**Fig 4.**
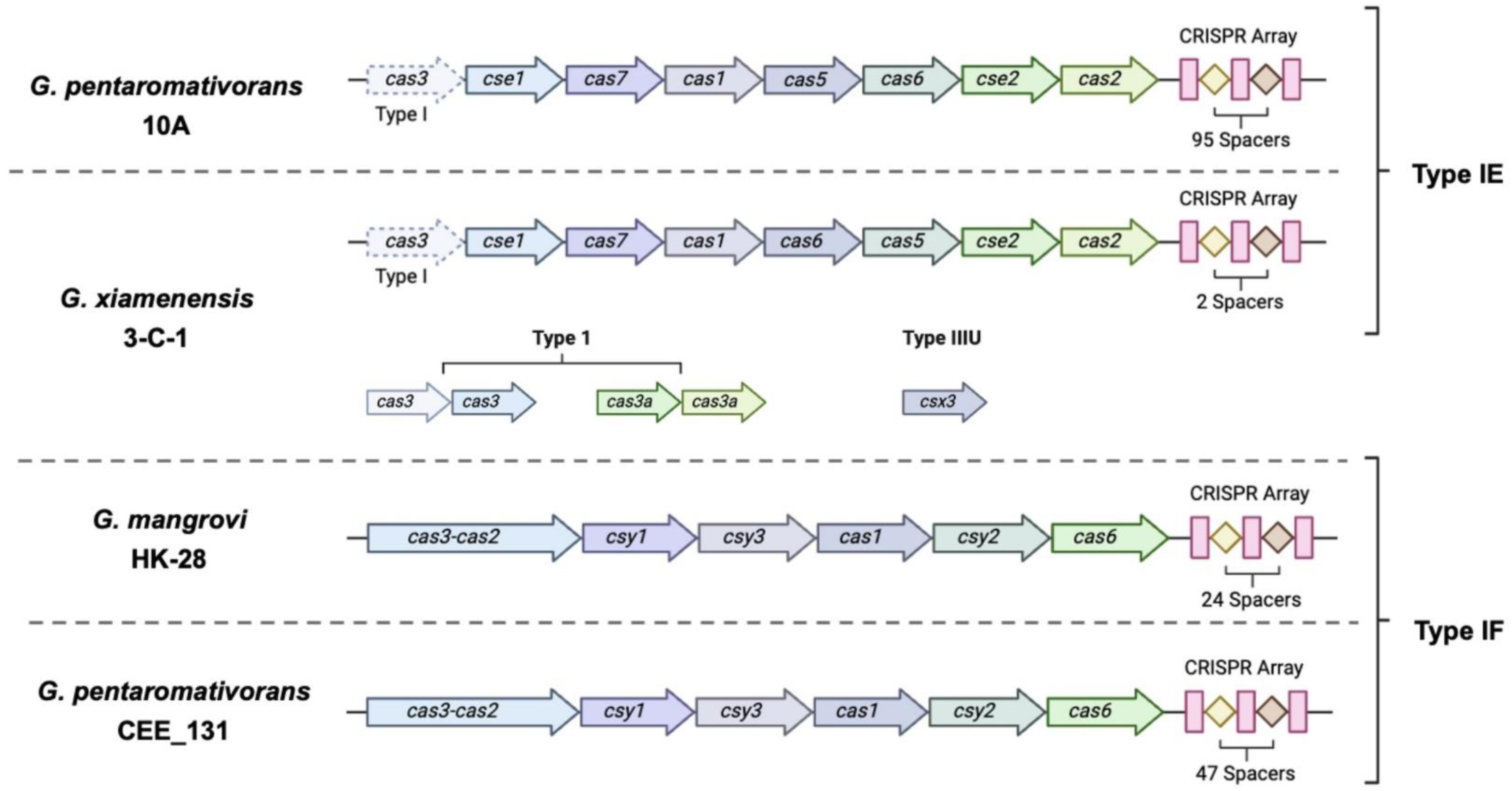
**Detection of CRISPR-Cas systems in *Gallaecimonas* spp.** Operon organization of different CRISPR-Cas systems detected in genomes of *Gallaecimonas* spp., annotated with the clustered regularly interspaced short palindromic repeats (CRISPR), spacers, and CRISPR-associated (Cas) proteins. Annotations are based on sequence similarities to known Cas proteins using HMM protein profiles and identified using CRISPRFinder.

### Global distribution of *Gallaecimonas*

Examination of the Global Biodiversity Information Facility (GBIF) 16S rRNA gene database [33] revealed sequences for *Gallaecimonas* spp. from a wide range of habitats ranging from polar regions to equatorial zones (Fig 5; S4 Table). In total, 536 16S rRNA gene OTUs assigned to *Gallaecimonas* were detected in the database. Of these 536 OTUs, 37.18% were resolved to species, with *G. pentaromativorans* representing 0.42% and *G. xiamenensis* representing 36.76% of the total. The remaining 62.82% of sequences were assigned to other *Gallaecimonas* spp., including *G. kandeliae*, *G. mangrovi*, and unclassified *Gallaecimonas,* including *Gallaecimonas sp*. SSL4-1, *Gallaecimonas sp*. MA-8, and *Gallaecimonas sp*. L2 (Fig 5A; S4 Table). Notably, most *Gallaecimonas*-containing samples were from marine environments (74.83%), including the pelagic ocean, saltmarshes, beaches, coral reefs, aquaculture ponds, and tidal flats (Fig 5B; S4 Table). Nonetheless, the footprint of *Gallaecimonas* also extends to other environments, including sediments (9.33%), forest soil (1.87%), estuaries (0.19%), rivers (0.19%), and polar systems (0.19%), with the remainder to be classified (13.41%) (Fig 5B). Overall, the relative abundance of *Gallaecimonas* 16S rRNA gene sequences in these samples (n = 536) ranged between 0.000048 and 3.99% (Fig 5A; S4 Table).

**Fig 5.**
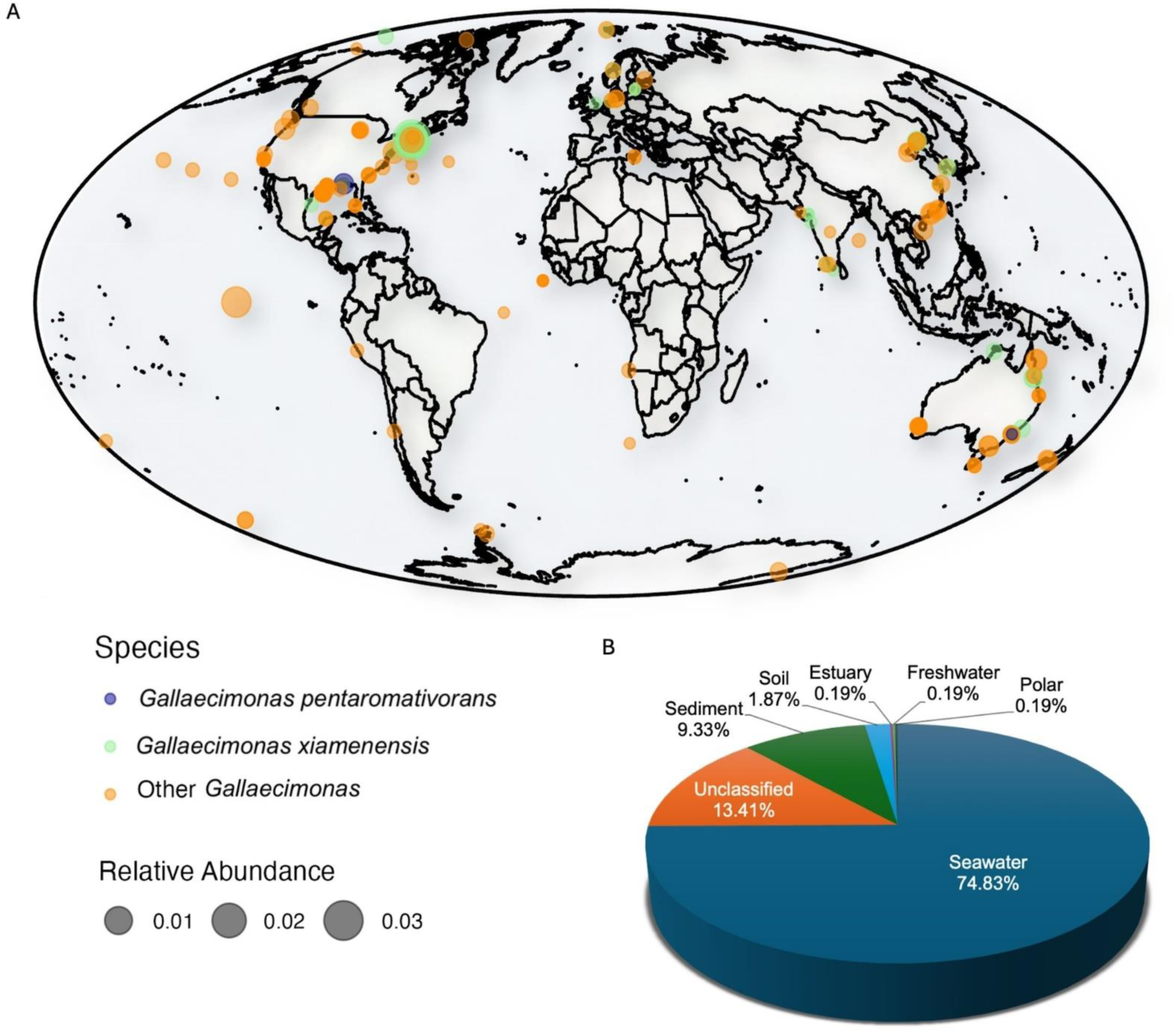
Distribution of *Gallaecimonas* across locations and habitats. (A) The global distribution of 16S rRNA gene sequences assigned to *Gallaecimonas* spp. was assessed using 536 sequences sourced from the GBIF Database. Orange and green indicate sequences assigned to *G. pentaromativorans* and *G. xiamenensis*, respectively, while “Other *Gallaecimonas*” comprises *G. kandeliae*, *G. mangrove*, and unclassified *Gallaecimonas* spp. The circle size represents the relative abundance of sequences assigned to *Gallaecimonas* spp. within the prokaryotic community. (B) The proportion of 16S rRNA gene sequences assigned to *Gallaecimonas* spp. across environments. Seawater, estuary, and freshwater correspond to water samples from the respective environments. Polar represents the cryosphere, and sediment includes samples from marine (e.g., pelagic sediment cores, benthic continental shelf), rivers, wetlands, saltmarshes, and tidal mudflats.

## Discussion

Bacteria in the genus *Gallaecimonas* have important industrial and ecological roles as biosurfactant producers and oil degraders [1, 2]. Here, we report on *G. pentaromativorans* strain 10A, the first isolate of *Gallaecimonas* spp. reported from an animal host, the Pacific oyster (*Magallana gigas*, a.k.a. *Crassostrea gigas*). The sequencing of strain 10A provides the first complete genome for *G. pentaromativorans*, adding to our repertoire of genomic knowledge for this important genus. This discovery, in addition to the genomic and biogeographic analysis of *Gallaecimonas*, enriches our knowledge of the ecological and functional roles of *Gallaecimonas*.

### Strain 10A has genetic potential for PAH and xenobiotic degradation

PAHs are naturally and anthropogenically produced, ubiquitous, recalcitrant, and have a high potential for bioaccumulation and carcinogenesis [37, 38]. As PAHs and xenobiotic environmental pollutants are highly persistent, the ability to degrade these pollutants is particularly valued [38, 39]. *Gallaecimonas pentaromativorans* strain 10A possesses genes for degrading xenobiotic compounds (ethylbenzene, caprolactam, chloroalkanes, chloroalkenes, atrazine, styrene, nicotinate, and nicotinamide) and PAHs (benzoate and naphthalene), suggesting it may have broad applications in bioremediation efforts (S3 Fig; Fig. 4C). This is consistent with the ability of *G. pentaromativorans* strain CEE_131 to break down PAHs like pyrene and benzo[a]pyrene [1]. While *G. xiamenensis* lacks this ability, *G. mangrovi* and *G. kandeliae* exhibit varying degrees of hydrocarbon degradation potential [4, 5]. While, experimental validation is needed to confirm its full potential, the abundance of aldehyde dehydrogenase and cytochrome P450 CDS in strain 10A suggests the ability to degrade these substances. More specifically, PAHs can induce their own metabolism by upregulating xenobiotic-metabolizing enzymes, such as cytochrome P450, by activating cellular receptors [40]. The predicted metabolic versatility of strain 10A showcases its potential as an effective bioremediation agent for contaminated environments [38, 40].

### Strain 10A has potential for PAH detoxification in oysters

The isolation of *G. pentaromativorans* strain 10A from a Pacific oyster hints at the possible presence of PAHs. Oysters, due to their filter-feeding behavior and stationary lifestyle, are prone to accumulate PAHs, pollutants that are pervasive in marine environments due to anthropogenic activities [39, 41, 42]. Thus, PAHs can accumulate in oysters and reach levels higher than in the surrounding waters, highlighting their susceptibility to environmental pollutants [42–44]. Consequently, the presence of *Gallaecimonas* in oysters suggests a role in detoxifying PAHs, as well as being a potential bioindicator for PAH contamination [42].

### Strain 10A is a candidate for biosurfactant producer

Strain 10A is predicted to encode a pathway consisting of (i) a two-component system, (ii) quorum sensing, (iii) biofilm formation, and (iv) flagellar assembly that was not found in the genomes of other isolates of *Gallaecimonas* spp. This pathway is hypothesized to be a biosurfactant-synthesis pathway, which typically encodes for processes such as attachment and dissociation of cells to surfaces, motility, quorum sensing, and biofilm formation [45]. More specifically, the two-component system and quorum sensing provide mechanisms to coordinate actions for rapid responses to environmental cues triggering bacterial motility and biofilm development [46–49]. Biofilm formation and flagellar assembly enhance the capacity for colonizing diverse ecological niches. Collectively, the presence of these pathways is consistent with the potential for biosurfactant synthesis, which could have industrial applications for oil and food production, as well as bioremediation [50]. Their natural role in emulsifying hydrophobic substrates enhances the availability of nutrients necessary for microbial growth, further contributing to their ecological importance [45, 51]. In general, isolates of *Gallaecimonas* are one of the most efficient biosurfactant-producing bacteria [2]. Strain 10A adds to this repertoire, and potentially offers another system for biosurfactant production, and a sustainable alternative to chemical surfactants using renewable substrates and natural processes [45, 50, 51].

### Strain 10A has diverse metabolism versatility

The pan-genomic analysis of currently available *Gallaecimonas* genomes revealed that strain 10A shares with others a number of the core genes involved in pathways dealing with cellular and environmental information processing (Fig. 3; S1-2 Figs; S1-2 Tables). These common cellular pathways are also highly conserved pathways found in *Gammaproteobacteria* [52], encompassing critical functions such as cell growth and death, the citrate cycle, amino-acid metabolism, cell-wall metabolism, and the synthesis of peptidoglycan, a crucial component of bacterial cell walls [53]. Moreover, the conserved pathways for environmental information processing cover a wide array of functions vital for bacterial physiology and ecological interaction. For example, the osmotic stress response pathway, mediated by *MtrB-MtrA*, enables bacteria to regulate their internal osmolarity in response to fluctuating environmental conditions, which is particularly valuable given the presence of the genus in estuaries and intertidal environments [54]. Similarly, the regulation of membrane lipid fluidity by *DesK-DesR* is critical for preserving membrane integrity and function, especially in response to temperature fluctuations [55]. These capacities allow strain 10A and other *Gallaecimonas* spp. to adapt to varying environmental conditions, such as salinity and temperature, and maintain cellular function, likely contributing to the widespread presence of members of this genus across diverse environments.

Also conserved within the genus are genes involved in capsule-synthesis pathways, including *ResC-ResD-ResB*, which play a vital role in providing protection against both host immune responses and environmental stresses [56]. However, capsules have not been reported in *Gallaecimonas*, suggesting that this pathway might have a different role or that the conditions needed for capsule formation have not yet been established. Interestingly, a conserved pathogenic cycle (map05111) found in both *Vibrio cholerae* and *Gallaecimonas* spp., suggests that both may be able to induce disease [57]. Ultimately, these conserved pathways highlight the sophisticated mechanisms that strain 10A and other *Gallaecimonas* may employ for adaptation and proliferation in diverse environments.

Like other *Gallaecimonas*, strain 10A also encodes a diverse array of proteins predicted to be involved in carbon, nitrogen, and sulfur cycling (Fig. 3; S1-2 Tables; S1-3 Figs). Notably, the predicted capacity for methanol oxidation, likely for the acquisition of carbon and energy, suggests involvement in carbon cycling within marine and soil ecosystems [58]. Moreover, genomic analysis revealed a complete nitrate-assimilation pathway and nitrate-assimilating genes (NAS), classifying strain 10A as a nitrate-assimilating bacteria (NAB) [58, 59]. Additionally, the genomic makeup of strain 10A showcases both complete assimilatory and dissimilatory sulfate-reduction pathways, indicating chemosynthetic capabilities [60]. Sulfate-reducing bacteria (SRB) are important in sustaining the diversity and stability of marine bacterial communities [61]. Thus, strain 10A is predicted to have the capacity to play a key role in nitrogen, carbon, and sulfur cycling across a diverse range of environments.

### Strain 10A has *higBA* toxin/anti-toxin system to deal with stress response

Strain 10A is predicted to encode a *higBA* type II toxin-antitoxin (TA) system that is not found in other isolates of *Gallaecimonas* (Fig. 3C; S1-2 Tables). The system, consisting of *HigB-1* and *HigA-1*, responds to stress conditions, and serves as a potential defense mechanism against stressors (e.g., cleaving foreign mRNA and encoding bacteriostatic toxins) [62]. Additionally, the TA system helps with adaptation to environmental challenges. For instance, in other gram-negative bacteria, such as *Pseudomonas aeruginosa,* the system regulates swarming and biofilm formation [62, 63], and has been linked to survival strategies in fluctuating environmental conditions [62].

### CRISPR-Cas system was detected in *G. pentaromativorans* strain 10A, and widespread in *Gallaecimonas*

CRISPR-Cas systems, which function as adaptive prokaryotic defense mechanisms against foreign plasmids and viruses, were detected in *G. pentaromativorans* strain 10A [36] (Fig. 4; S8 Fig; S3 Table). This discovery is particularly notable, as CRISPR-Cas systems had not been previously reported in *Gallaecimonas*. Upon examination of genomic data from all five currently available *Gallaecimonas* isolates, including strain 10A, we found that CRISPR-Cas systems are widespread in *Gallaecimonas*, with four out of the five genomes containing both CRISPR arrays and Cas clusters (Fig. 4; S4-8 Figs). These spanned three different species (*G. pentaromativorans*, *G. mangrovi*, and *G. xiamenensis*) within the genus (Fig. 4; S5-8 Figs); only *G. kandeliae* did not exhibit a full CRISPR-Cas system but with merely CRISPR arrays detected (S4 Fig). The CRISPR-Cas cassettes found in *Gallaecimonas* belong to class 1 Type IE and IF (Fig. 4), which are common in gram-negative bacteria [64]. The Type IE systems in *G. pentaromativorans* strain 10A and *G. xiamenensis* strain 3-C-1 have a cascade, a multiprotein surveillance complex [64]. The Type IF system seen in *G. mangrovi* strain HK-28 and *G. pentaromativorans* strains CEE_131 (DSM21945) have a *Csy* complex with *cas2* and *cas3* genes fused into a single open reading frame [64]. Strikingly, strain 10A has one full Type IE CRISPR-Cas cassette with 95 spacers (Fig. 4; S8 Fig; S3 Table), indicating historical and frequent encounters with foreign genetic elements, such as viruses.

### Strain 10A is morphologically similar to other *Gallaecimonas*

Similar to other isolates of *Gallaecimonas* spp., *G. pentaromativorans* strain 10A is rod-shaped, with cells about 0.5-μm wide and 2-μm long, although other members of the genus have been reported to range from 0.3- to 0.9-μm in width and 1- to 3-μm in length [3–5]. Based on the soft agar stab test, strain 10A appears to be motile, while preliminary TEM observations hint at the presence of a polar flagellum akin to that reported for *G. pentaromativorans* strain CEE_131 [1]; whereas, *G. xiamenensis* has an amphitrichous arrangement [3], and no flagella have been reported for *G. mangrove* [4] or *G. kandeliae* [5]. Furthermore, we identified the presence of genes (e.g., *PilB-PilV*) involved in type IV fimbriae (also known as type IV pili) synthesis in *G. pentaromativorans* strain 10A (S1 Table). This finding suggests that strain 10A may exhibit ’twitching motility’, a process facilitated by the extension of long, thin fimbriae from the cell wall, which are involved in surface adherence and movement [65–67]. Type IV fimbriae are multifunctional structures and may play a variety of roles, including host attachment, biofilm formation, and/or potentially pathogenicity [67–71]. Interestingly, some of these type-IV-fimbriae-associated genes (e.g., *PilS-PilR*) were also identified as core genes shared among all four *Gallaecimonas* species described to date, highlighting their potentially important role within the genus.

*Gallaecimonas* colonies, including strain 10A, are consistently characterized as being smooth, circular, convex, colorless to gray colored, and 0.1- to 3-mm in diameter [1, 3–5].

### *Gallaecimonas* are widespread across environments

Bacteria in the genus *Gallaecimona*s demonstrate remarkable ecological versatility, as evidenced by their widespread presence across different environments at all latitudes (Fig 5; S4 Table). Members of the genus have primarily been reported from marine environments, aligning with isolations from a crude oil-degrading consortium in seawater [3], mangrove sediments [4, 5], and intertidal sediments [1]. However, environmental sequencing data indicate that their range also extends to soils, estuaries, rivers, and the cryosphere (Fig 5; S4 Table). Thus, bacteria in the genus exist across a wide range of salinities, temperatures, and environmental conditions. In fact, members of the genus have been grown at 10 to 45 °C, pH 5 to 10, and NaCl concentrations from 0 to12 % [1, 3–5].

The species-specific biogeography of *Gallaecimonas* is limited in the current investigation because the GBIF 16S rRNA gene database only includes samples from 2003 to 2019, which does not reflect recent taxonomic updates and overlooks species such as *G. mangrovi* and *G. kandeliae*. Similarly, *G. pentaromativorans*, including strain 10A, was only detected on two occasions in marine sediment and seawater. Such a low occurrence of *G. pentaromativorans* in natural environments might reflect its host-associated nature (e.g., strain 10A), suggesting that we need to search for them in animal microbiomes (e.g., oysters). Moreover, the distribution of *G. pentaromativorans* strain 10A in global ecosystems also requires more sampling efforts and accurate taxonomy assignment utilizing the most current 16S SSU database (e.g., the future database with strain 10A included).

## Conclusion

Here, we present the first complete genome published for the species, *Gallaecimonas pentaromativorans*. The isolation and genomic characterization of *G. pentaromativorans* strain 10A from Pacific oyster, alongside comparative genomics with other isolates, demonstrates the remarkable genomic potential of strain 10A, and of members of the genus, more broadly. As well, by mining environmental data, we show the widespread distribution of the genus globally and across environments (e.g., seawater, sediment, and oysters). Functionally, *G. pentaromativorans* strain 10A shows its potential to be a biosurfactant producer and a key player in critical environmental processes, including carbon cycling and the metabolism of sulfur and nitrogen, acting as both a nitrate-reducing and sulfate-reducing bacterium. Notably, *G. pentaromativorans* has the potential to be an important bioremediation agent through PAH and xenobiotic compound degradation and metabolism. Given that *Gallaecimonas* has been proposed as a bio-indicator for PAHs, the isolation of strain 10A from oysters suggests the possibility of PAH contamination and degradation in oysters.

## Data availability

The genome sequence of *Gallaecimonas pentaromativorans* strain 10A is deposited in Genbank under accession number CP152402. Raw sequencing data can be accessed through NCBI Sequence Read Archive (SRA) with accession numbers SRR28797806 to SRR28797807.

## Acknowledgments

We respectfully acknowledge that this research was conducted on the unceded traditional territories of the xʷməθkʷə^y̓^ əm (Musqueam), Sḵwx̱wú7mesh (Squamish), and səlilwətaɬ (Tsleil- Waututh) Nations. We appreciate the assistance of RKS Labs in obtaining the oyster samples; and thank C.J. Huang and M. Daspe for assistance with culturing, Q. Yang for assistance with TEM sample preparation, and staff members at the UBC BioImaging Facility (RRID:SCR_021304) for technical assistance.

## Author Contribution

A.M.C. and C.A.S. acquired the oyster samples. A.M.C. directed the bacterial isolation. A.W. and A.M.C. isolated and cultured *G. pentaromativorans* strain 10A. A.M.C. performed microscopy and analysis. Y.G. and K.X.Z. conducted the bioinformatic analysis and interpreted the results. Y.G. wrote the draft manuscript, which was revised by K.X.Z., C.A.S. and A.M.C.

## Supplementary Information

### 1. Supplementary Table Legends

**Table S1. Functional annotation of genes in *Gallaecimonas pentaromativorans* strain 10A**. Data consists of predicted seed ortholog (i.e., gene family), e-value, score, eggNOG OGs, max annotation level, COG category, description, preferred name, GOs, EC, KEGG profile data (ko, Pathway, Module, Reaction, rclass, TC), BRITE, CAZy, BiGG Reaction, and PFAMs (i.e., protein family) using eggNOG-mapper.

**Table S2. Metadata for comparative genomics.** The eggNOG-mapper generated data of (i) common KEGG Ortholog (ko) profiles for *Gallaecimonas* spp., (ii) common ko profiles for *G.pentaromativorans*, and (iii-vii) unique ko profiles for each *Gallaecimonas* spp.

Table S3. The clustered regularly interspaced short palindromic repeats (CRISPR) and CRISPR spacer sequences detected in the genome of *G. pentaromativorans* strain 10A.

**Table S4. Metadata for analysis of the distribution of *Gallaecimonas* across geolocations and habitats.** Data sourced from the GBIF Database and cleaned to include gbifID, dynamicProperties, occurrenceID, materialSampleID, eventid, sampleSizeValue, sampleSizeUnit, continent, waterbody, decimalLatitude, decimalLongitude, genus, specificEpithet, depth, Scientific Name.

### 2. Supplementary Figures

**Fig S1.**
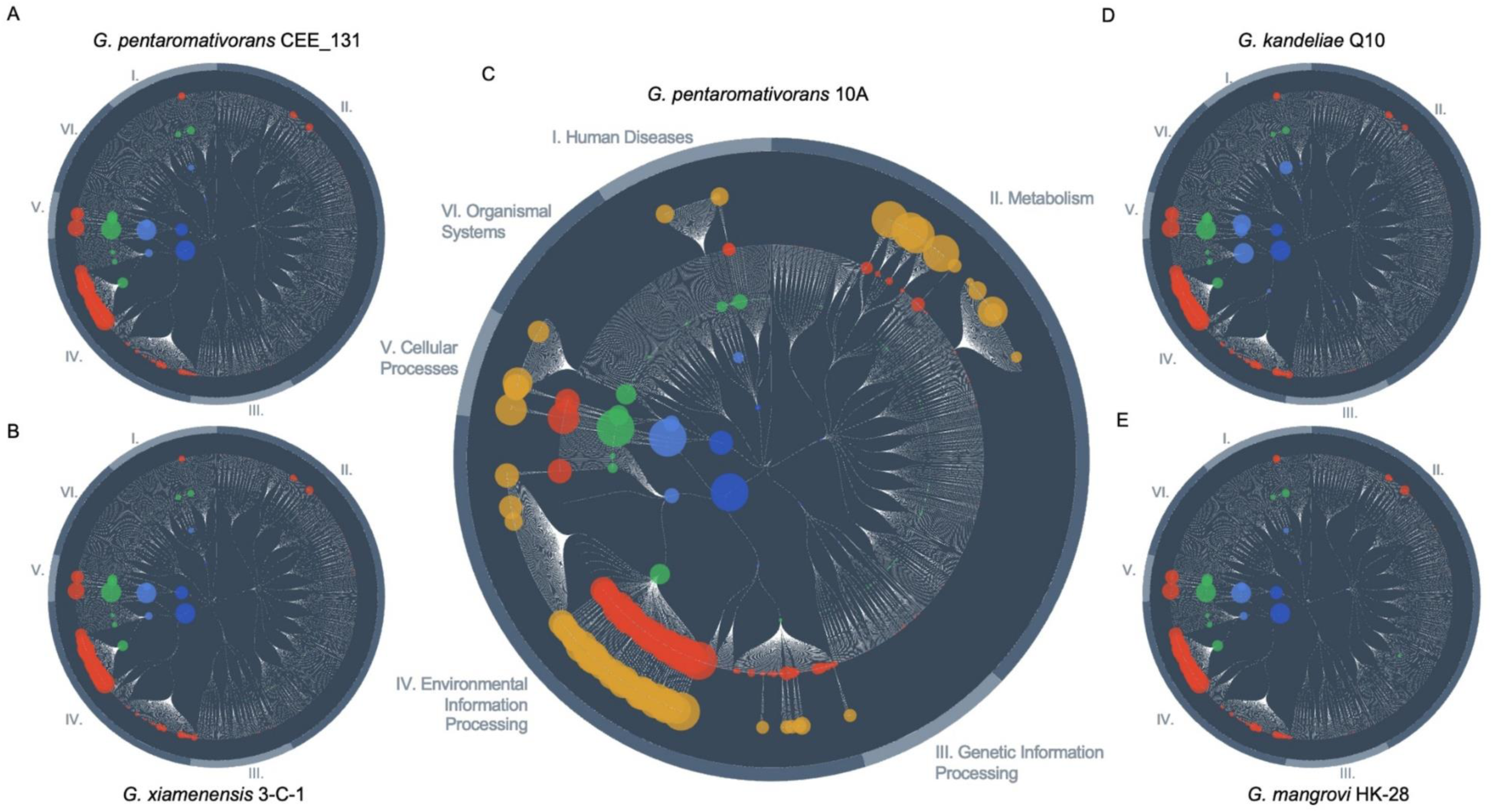
**Functional maps of *Gallaecimonas.*** Functional genomic maps for the five isolates of *Gallaecimonas* spp. for which full genome sequences are available (A) *G. pentaromativorans* strain CEE_131 (Leibniz Institute DSMZ culture collection ID: DSM21945); (B) *G. xiamenensis* strain 3-C-1; (C) *G. pentaromativorans* strain 10A; (D) *G. kandeliae* strain Q10; (E) *G. mangrovi* strain HK-28) made with FuncTree v.0.8.4. Node color is depicted as follows from the outermost to innermost rings of the map: yellow (only for strain 10A) denotes KEGG Orthology (KO); red signifies KEGG Module; green indicates KEGG Pathways; light blue represents biological processes; dark blue represents biological categories. The position on the circle represents category: I. Human Diseases, II. Metabolism, III. Genetic Information Processing, IV. Environmental Information Processing, V. Cellular Processes, VI. Organismal Systems. The node size corresponds to the value of the standard deviation of the KO’s relative abundance assigned to that function.

**Fig S2.**
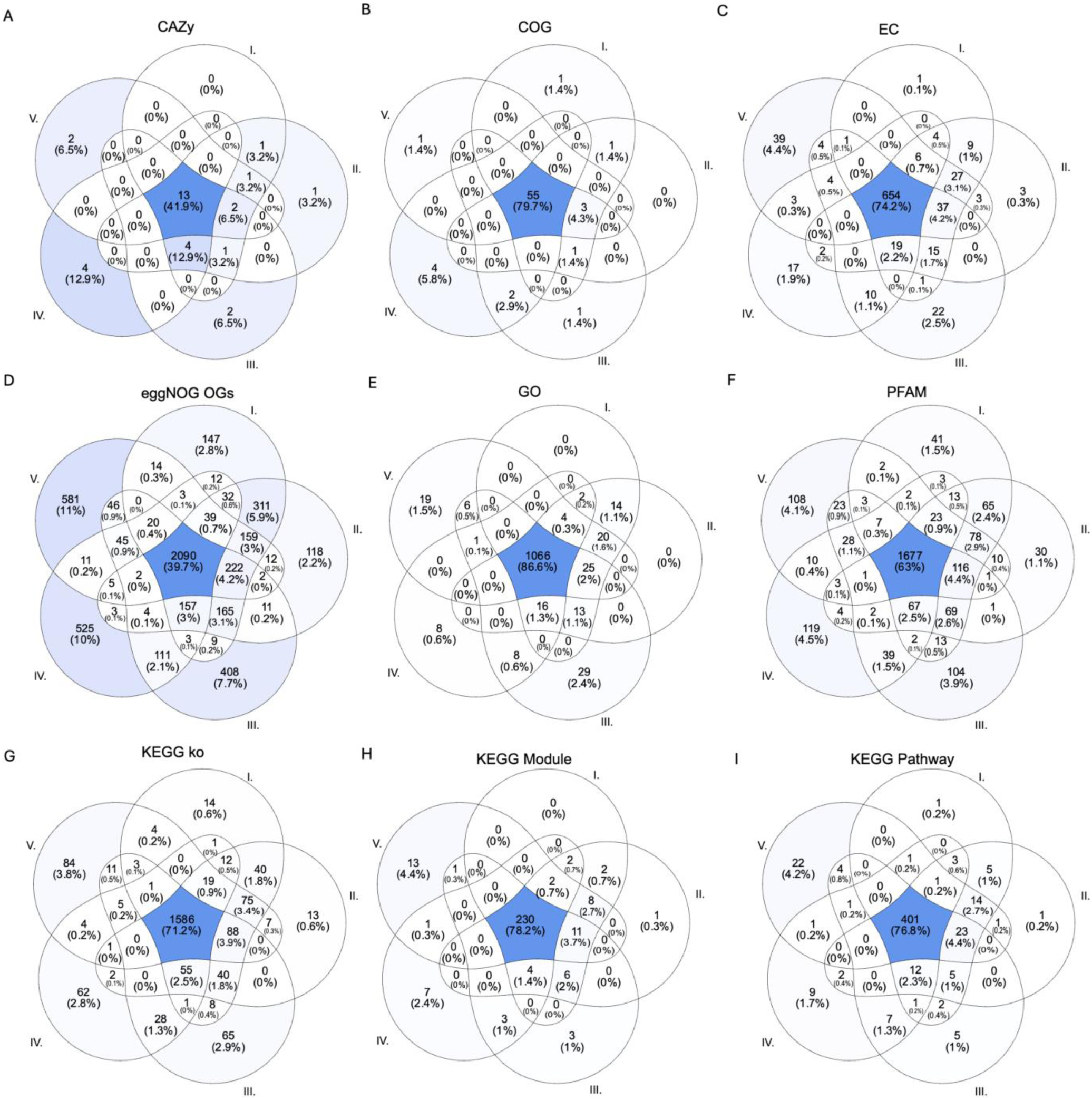
**Comparative genomic features of *Gallaecimonas* spp.** Venn diagrams illustrating the distribution of (A) CAZy, (B) COG profiles, (C) EC ids, (D) eggNOG Ogs, (E) GO terms, (F) PFAMs (i.e., protein family), (G) KEGG ko profiles, (H) KEGG Module profiles, (I) KEGG Pathway profiles among genomes of *Gallaecimonas* spp. (I. *G. pentaromativorans* strain 10A; II. *G. pentaromativorans* strain CEE_131 (Leibniz Institute DSMZ culture collection ID: DSM21945); III. *G. xiamenensis* strain 3-C-1; IV. *G. kandeliae* strain Q10; V. *G. mangrovi* strain HK-28) according to eggNOG-mapper.

**Fig S3.**
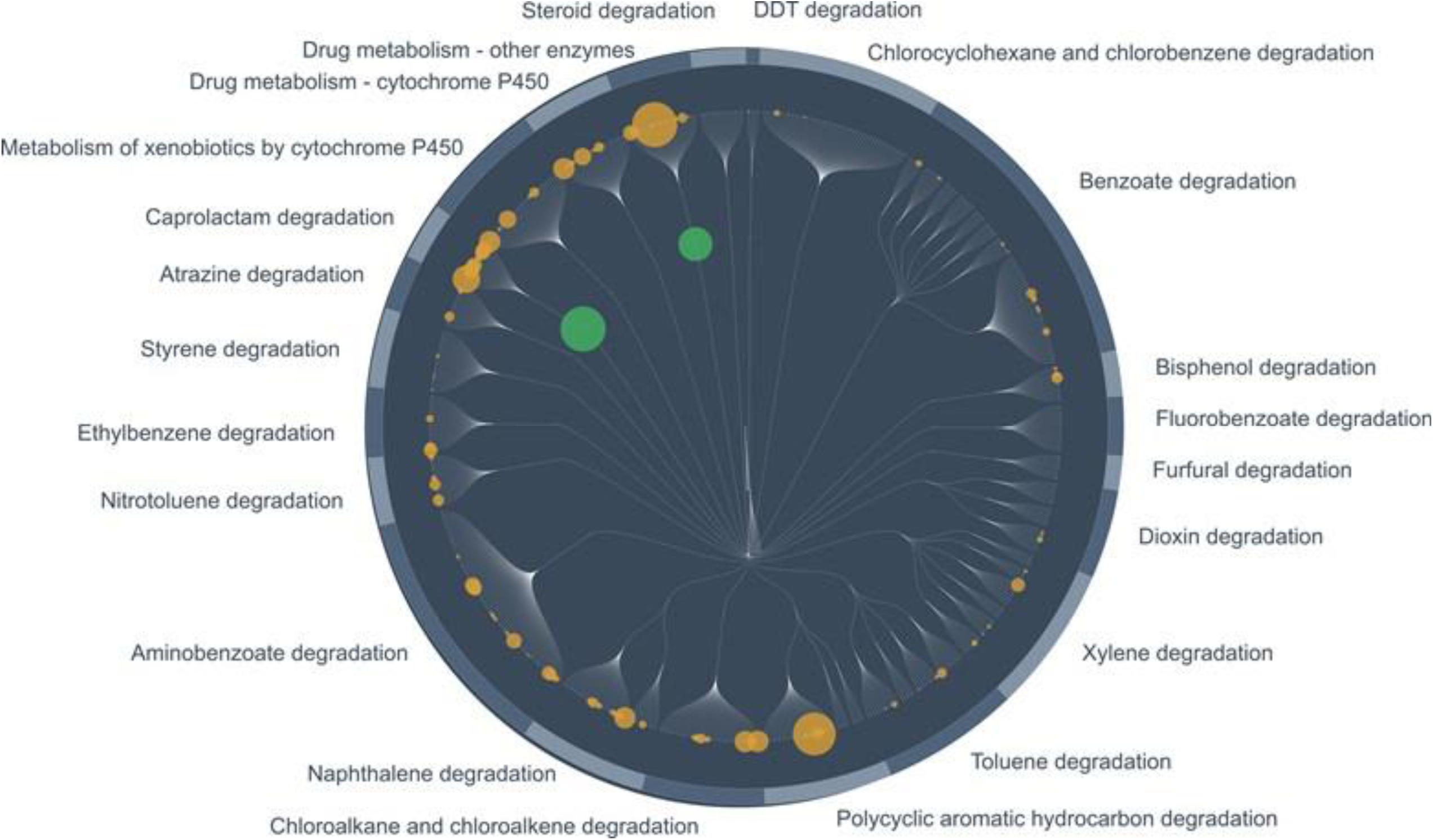
PAHs and xenobiotics degradation potential of *G. pentaromativorans* strain 10A. Functional map of the PAHs and xenobiotics degradation KO profiles with over 50% coverage for module and pathway made with FuncTree v.0.8.4. The outermost to innermost rings are as follows: yellow represents KEGG Orthology, and green represents KEGG Pathways. Node size corresponds to the value of the standard deviation of the KO’s relative abundance assigned to that function.

**Fig S4.**
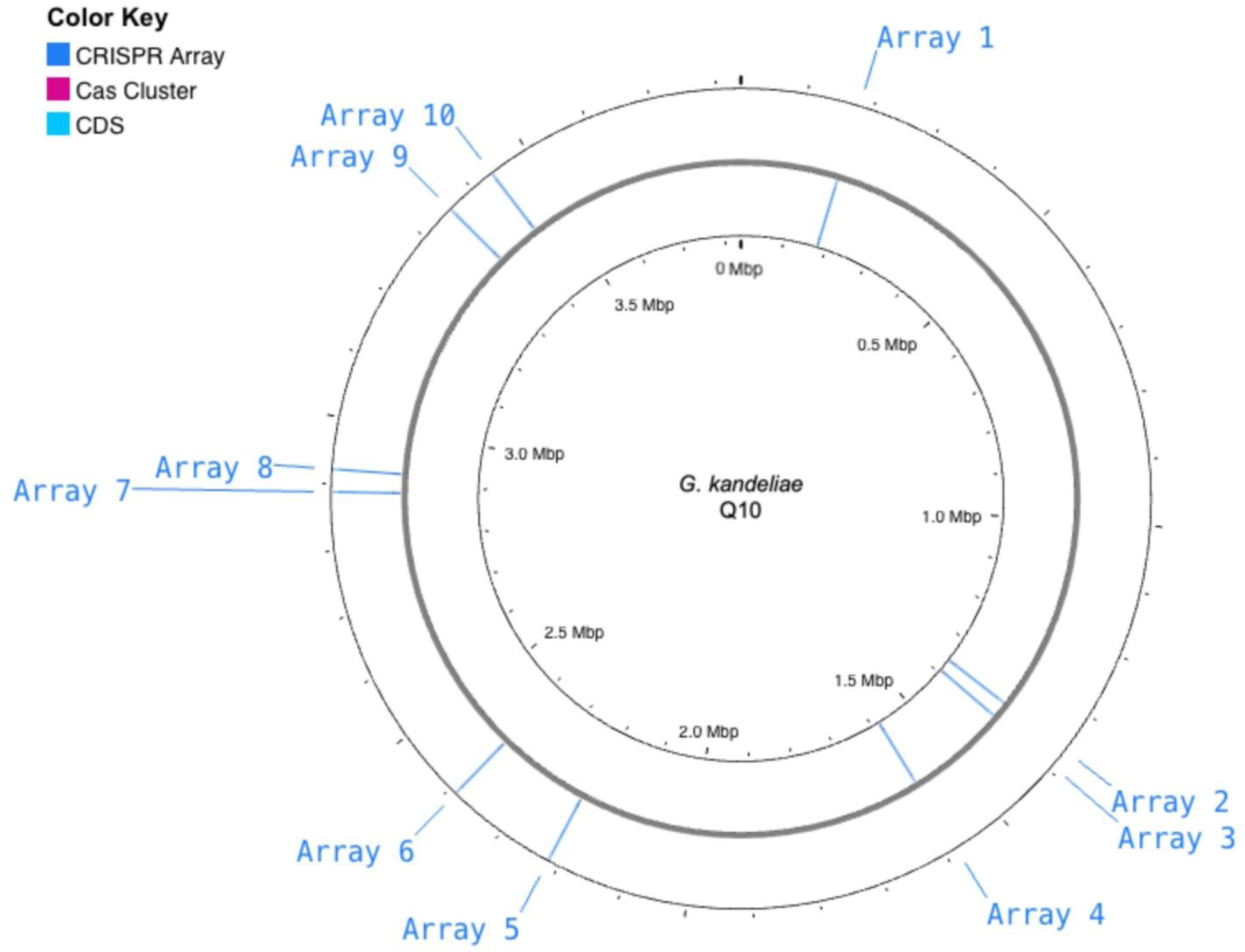
**CRISPR-Cas analysis of *G. kandeliae* strain Q10.** Genomic map of *G. kandeliae* strain Q10, annotated with the clustered regularly interspaced short palindromic repeats and spacers (CRISPR arrays), CRISPR-associated (Cas) proteins and clusters, and putative coding sequences (CDSs). Annotations are based on sequence similarities to known Cas proteins using HMM protein profiles and identified using CRISPRFinder.

**Fig S5.**
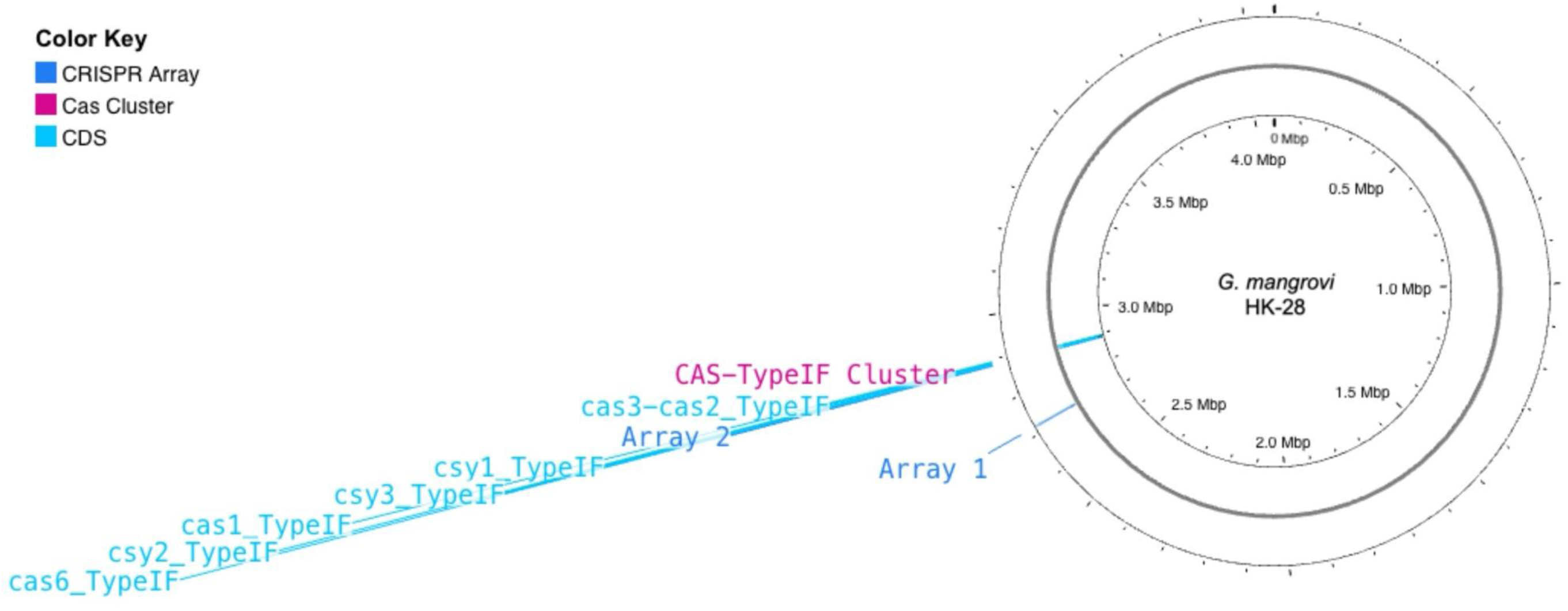
**CRISPR-Cas analysis of *G. mangrovi* strain HK-28.** Genomic map of *G. mangrovi* strain HK-28, annotated with the clustered regularly interspaced short palindromic repeats and spacers (CRISPR arrays), CRISPR-associated (Cas) proteins and clusters, and putative coding sequences (CDSs). Annotations are based on sequence similarities to known Cas proteins using HMM protein profiles and identified using CRISPRFinder.

**Fig S6.**
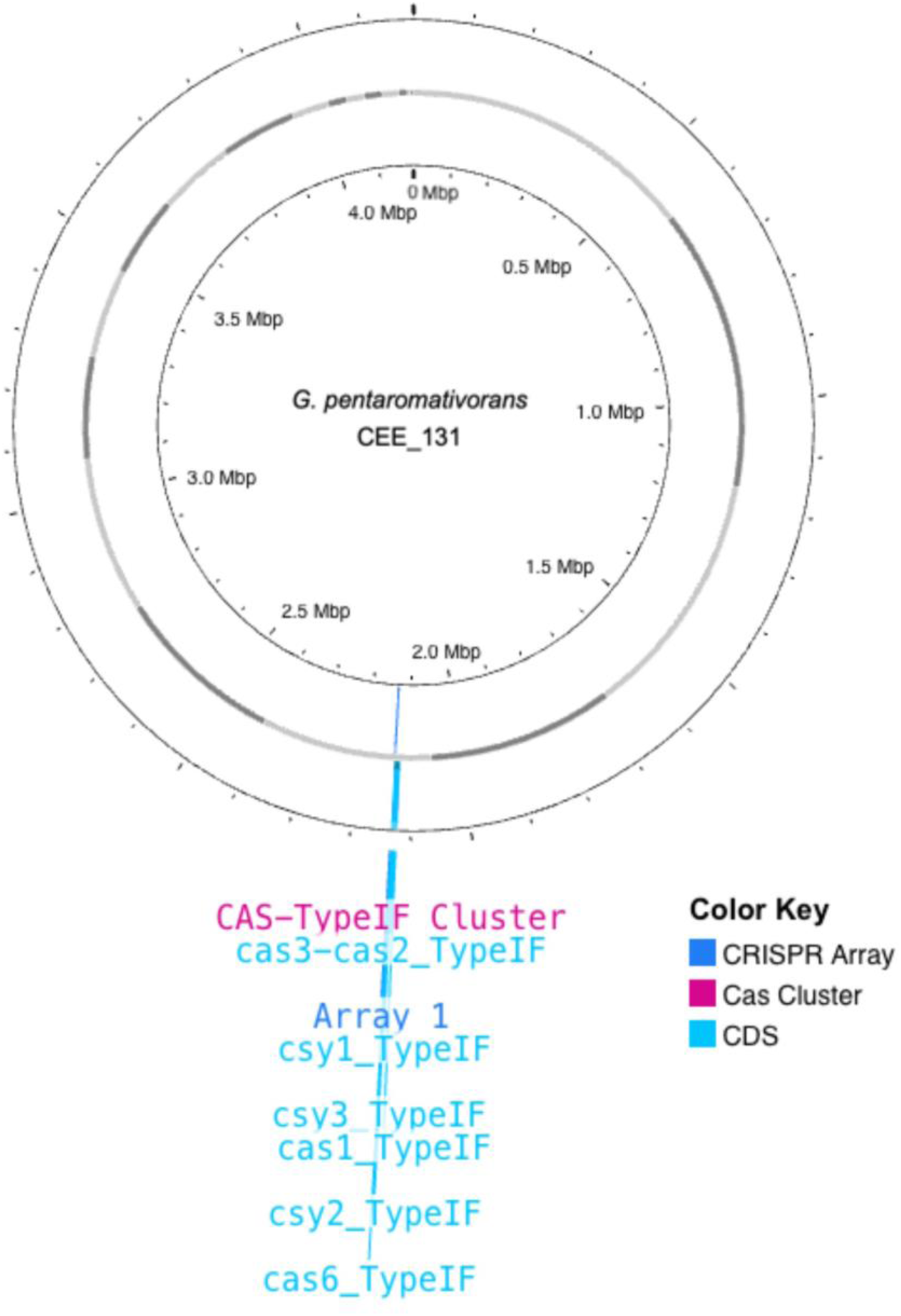
**CRISPR-Cas analysis of *G. pentaromativorans* strain CEE_131.** Genomic map of *G. pentaromativorans* strain CEE_131 (Leibniz Institute DSMZ culture collection ID: DSM21945), annotated with the clustered regularly interspaced short palindromic repeats and spacers (CRISPR arrays), CRISPR-associated (Cas) proteins and clusters, and putative coding sequences (CDSs). Annotations are based on sequence similarities to known Cas proteins using HMM protein profiles and identified using CRISPRFinder.

**Fig S7.**
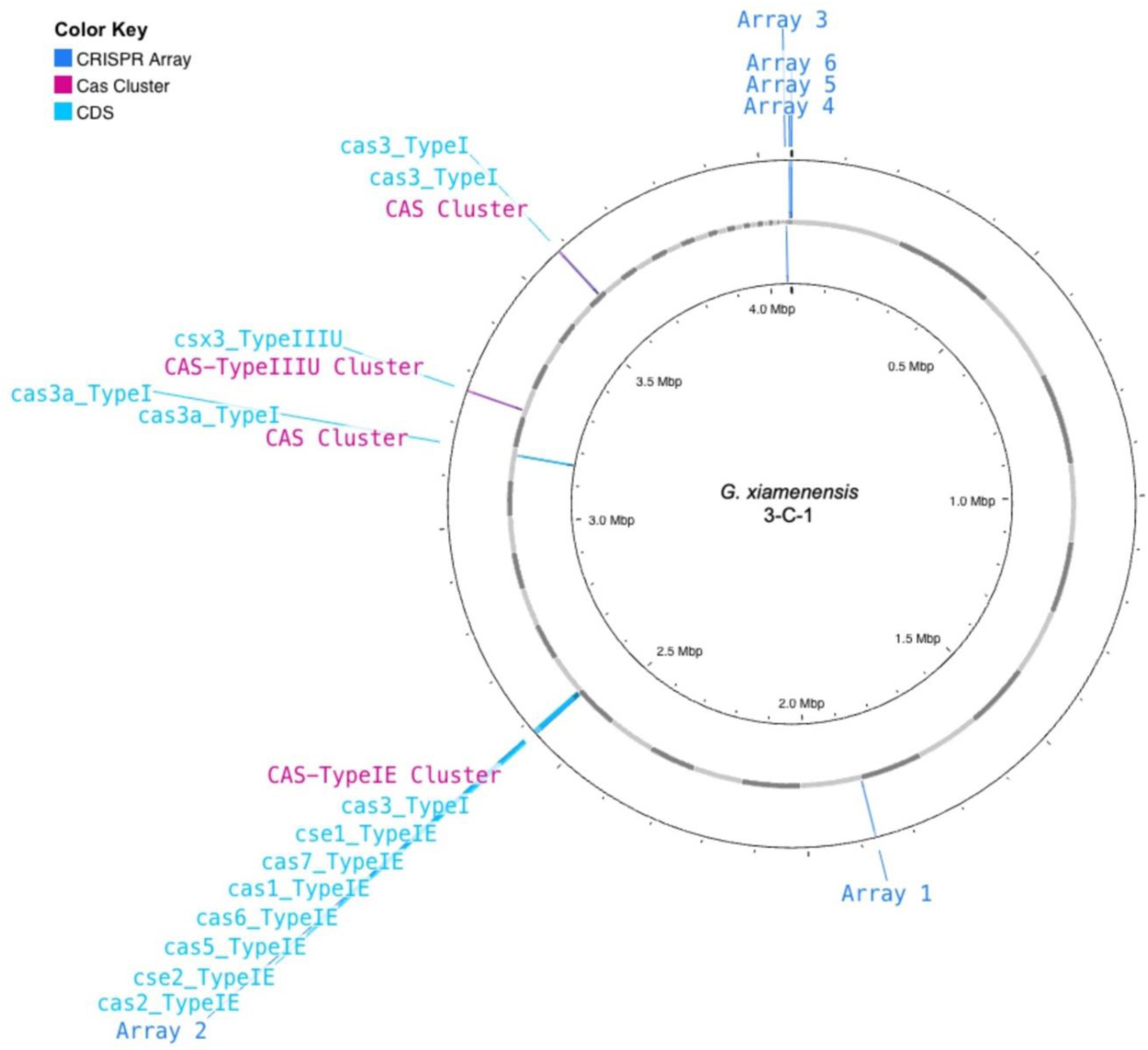
**CRISPR-Cas analysis of *G. xiamenensis* strain 3-C-1.** Genomic map of *G. xiamenensis* strain 3-C01, annotated with the clustered regularly interspaced short palindromic repeats and spacers (CRISPR arrays), CRISPR-associated (Cas) proteins and clusters, and putative coding sequences (CDSs). Annotations are based on sequence similarities to known Cas proteins using HMM protein profiles and identified using CRISPRFinder.

**Fig S8.**
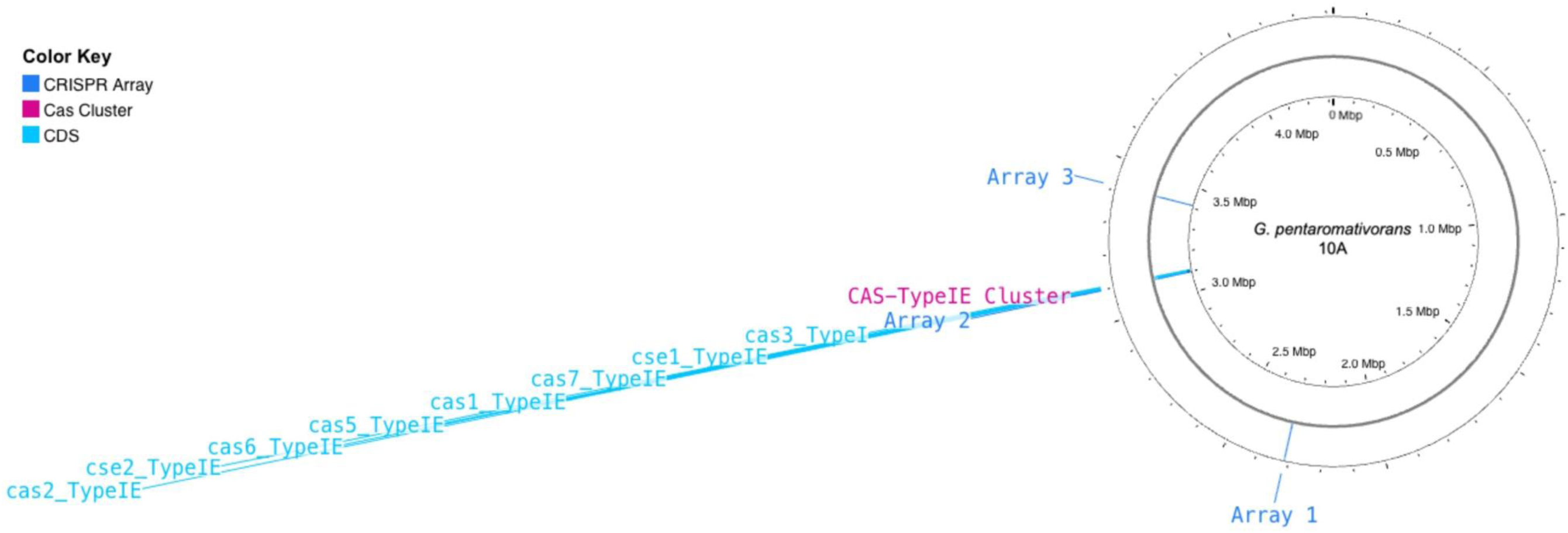
**CRISPR-Cas analysis of *G. pentaromativorans* strain 10A.** Genomic map of *G. pentaromativorans* strain 10A, annotated with the clustered regularly interspaced short palindromic repeats and spacers (CRISPR arrays), CRISPR-associated (Cas) proteins and clusters, and putative coding sequences (CDSs). Annotations are based on sequence similarities to known Cas proteins using HMM protein profiles and identified using CRISPRFinder.

## References

1. Rodriguez-Blanco A, Vetion G, Escande ML, Delille D, Ghiglione JF. *Gallaecimonas pentaromativorans* gen. nov., sp. nov., a bacterium carrying 16S rRNA gene heterogeneity and able to degrade high-molecular-mass polycyclic aromatic hydrocarbons. Int J Syst Evol Microbiol. 2010; 60:504-9. 10.1099/ijs.0.013532-0. PubMed PMID: 19654338.

2. Hassanshahian M. Isolation and characterization of biosurfactant producing bacteria from Persian Gulf (Bushehr provenance). Mar Pollut Bull. 2014; 86:361–6. 10.1016/j.marpolbul.2014.06.043. PubMed PMID: 25037876.

3. Wang J, Lai Q, Duan X, Fu Y, Wang L, Wang W, et al. *Gallaecimonas xiamenensis* sp. nov., isolated from seawater. Int J Syst Evol Microbiol. 2013; 63:930–3. 10.1099/ijs.0.042283-0. PubMed PMID: 22659502.

4. Zhang WY, Yuan Y, Su DQ, He XP, Han SB, Epstein SS, et al. *Gallaecimonas mangrovi* sp. nov., a novel bacterium isolated from mangrove sediment. Antonie Van Leeuwenhoek. 2018; 111:1855–62. 10.1007/s10482-018-1076-y.

5. Long M, Tang S, Fan H, Gan Z, Xia H, Lu Y. Description and genomic characterization of *Gallaecimonas kandeliae* sp. nov., isolated from the sediments of mangrove plant Kandelia obovate. Antonie Van Leeuwenhoek. 2023; 116:893–905. 10.1007/s10482-023-01851-y. PubMed PMID: 37358702.

6. Ding L, Xu P, Zhang W, Yuan Y, He X, Su D, et al. Three New Diketopiperazines from the Previously Uncultivable Marine Bacterium *Gallaecimonas mangrovi* HK-28 Cultivated by iChip. Chem Biodiversity. 2020; 17:e2000221. 10.1002/cbdv.202000221. PubMed PMID: 32347603.

7. Guillard RR, editor Culture of phytoplankton for feeding marine invertebrates. Culture of marine invertebrate animals: proceedings—1st conference on culture of marine invertebrate animals greenport; 1975: Springer. 10.1007/978-1-4615-8714-9_3.

8. . Suttle C, Chen F, Chan A, editors. Marine viruses: Decay rates, diversity and ecological implications. Int Mar Biotechnol Conf“IMBC-91” Develop Microbial Ser W Brown; 1992.

9. Suttle CA, Chen F. Mechanisms and rates of decay of marine viruses in seawater. Appl Environ Microbiol. 1992; 58:3721–9. 10.1128/aem.58.11.3721-3729.1992. PubMed PMID: 16348812.

10. Tittsler RP, Sandholzer LA. The use of semi-solid agar for the detection of bacterial motility. J Bacteriol. 1936; 31:575–80. 10.1128/jb.31.6.575-580.1936. PubMed PMID: 16559914.

1. Illumina. bcl2fastq Conversion Software 2019 [updated 2019]. Available from: https://support.illumina.com/sequencing/sequencing_software/bcl2fastq-conversion-software.html.

12. Wick R. Porechop: adapter trimmer for Oxford Nanopore reads 2018 [updated 2018]. Available from: https://github.com/rrwick/Porechop.

13. Wick RR, Judd LM, Gorrie CL, Holt KE. Unicycler: Resolving bacterial genome assemblies from short and long sequencing reads. PLoS Comput Biol. 2017; 13:e1005595. 10.1371/journal.pcbi.1005595. PubMed PMID: 28594827.

14. Parks DH, Imelfort M, Skennerton CT, Hugenholtz P, Tyson GW. CheckM: assessing the quality of microbial genomes recovered from isolates, single cells, and metagenomes. Genome Res. 2015; 25:1043–55. 10.1101/gr.186072.114. PubMed PMID: 25977477.

15. Menzel P, Ng KL, Krogh A. Fast and sensitive taxonomic classification for metagenomics with Kaiju. Nat Commun. 2016; 7:11257. 10.1038/ncomms11257. PubMed PMID: 27071849.

16. von Meijenfeldt FAB, Arkhipova K, Cambuy DD, Coutinho FH, Dutilh BE. Robust taxonomic classification of uncharted microbial sequences and bins with CAT and BAT. Genome Biol. 2019; 20:217. 10.1186/s13059-019-1817-x. PubMed PMID: 31640809.

17. Dong X, Strous M. An Integrated Pipeline for Annotation and Visualization of Metagenomic Contigs. Front Genet. 2019; 10:999. 10.3389/fgene.2019.00999. PubMed PMID: 31681429.

18. Li W, O’Neill KR, Haft DH, DiCuccio M, Chetvernin V, Badretdin A, et al. RefSeq: expanding the Prokaryotic Genome Annotation Pipeline reach with protein family model curation. Nucleic Acids Res. 2021; 49:D1020–D8. 10.1093/nar/gkaa1105. PubMed PMID: 33270901.

19. Chaumeil PA, Mussig AJ, Hugenholtz P, Parks DH. GTDB-Tk: a toolkit to classify genomes with the Genome Taxonomy Database. Bioinformatics. 2019; 36:1925–7. 10.1093/bioinformatics/btz848. PubMed PMID: 31730192.

20. Quast C, Pruesse E, Yilmaz P, Gerken J, Schweer T, Yarza P, et al. The SILVA ribosomal RNA gene database project: improved data processing and web-based tools. Nucleic Acids Res. 2013; 41:D590–6. 10.1093/nar/gks1219. PubMed PMID: 23193283.

21. Katoh K, Standley DM. MAFFT multiple sequence alignment software version 7: improvements in performance and usability. Mol Biol Evol. 2013; 30:772–80. 10.1093/molbev/mst010. PubMed PMID: 23329690.

22. Castresana J. Selection of conserved blocks from multiple alignments for their use in phylogenetic analysis. Mol Biol Evol. 2000; 17:540–52. 10.1093/oxfordjournals.molbev.a026334. PubMed PMID: 10742046.

23. Guindon S, Dufayard JF, Lefort V, Anisimova M, Hordijk W, Gascuel O. New algorithms and methods to estimate maximum-likelihood phylogenies: assessing the performance of PhyML 3.0. Syst Biol. 2010; 59:307–21. 10.1093/sysbio/syq010. PubMed PMID: 20525638.

24. Tamura K, Stecher G, Kumar S. MEGA11: Molecular Evolutionary Genetics Analysis Version 11. Mol Biol Evol. 2021; 38:3022–7. 10.1093/molbev/msab120. PubMed PMID: 33892491.

25. Cantalapiedra CP, Hernández-Plaza A, Letunic I, Bork P, Huerta-Cepas J. eggNOG- mapper v2: functional annotation, orthology assignments, and domain prediction at the metagenomic scale. Mol Biol Evol. 2021; 38:5825–9. 10.1093/molbev/msab293. PubMed PMID: 34597405.

26. Jin Z, Sato Y, Kawashima M, Kanehisa M. KEGG tools for classification and analysis of viral proteins. Protein Sci. 2023; 32:e4820. 10.1002/pro.4820. PubMed PMID: 37881892.

27. Grant JR, Enns E, Marinier E, Mandal A, Herman EK, Chen CY, et al. Proksee: in-depth characterization and visualization of bacterial genomes. Nucleic Acids Res. 2023; 51:W484–W92. 10.1093/nar/gkad326. PubMed PMID: 37140037.

28. Uchiyama T, Irie M, Mori H, Kurokawa K, Yamada T. FuncTree: Functional Analysis and Visualization for Large-Scale Omics Data. PLoS One. 2015; 10:e0126967. 10.1371/journal.pone.0126967. PubMed PMID: 25974630.

29. Grissa I, Vergnaud G, Pourcel C. CRISPRFinder: a web tool to identify clustered regularly interspaced short palindromic repeats. Nucleic Acids Res. 2007; 35:W52–7. 10.1093/nar/gkm360. PubMed PMID: 17537822.

30. Lai Q, Wang L, Wang W, Shao Z. Genome sequence of *Gallaecimonas xiamenensis* type strain 3-C-1. J Bacteriol. 2012; 194:6937. 10.1128/JB.01854-12. PMID: 23209203.

31. Wilkinson L. ggplot2: elegant graphics for data analysis by WICKHAM, H. Oxford University Press; 2011. 10.1007/978-0-387-98141-3.

32. Yan L, Yan ML. Package “ggvenn”. CRAN; 2021. https://cran.r-project.org/web/packages/ggvenn/readme/README.html.

33 Org G. Global biodiversity information facility. 2022. https://www.gbif.org/.

34. Pebesma EJ. Simple features for R: standardized support for spatial vector data. R J. 2018; 10:439. https://cran.r-project.org/web/packages/sf/index.html.

35. Brownrigg MR. Package ‘maps’. R package. 2013. https://cran.r-project.org/web/packages/maps/index.html.

36. Barrangou R, Fremaux C, Deveau H, Richards M, Boyaval P, Moineau S, et al. CRISPR provides acquired resistance against viruses in prokaryotes. Science. 2007; 315:1709–12. 10.1126/science.1138140. PubMed PMID: 17379808.

37. Haritash AK, Kaushik CP. Biodegradation aspects of polycyclic aromatic hydrocarbons (PAHs): a review. J Hazard Mater. 2009; 169:1–15. 10.1016/j.jhazmat.2009.03.137. PubMed PMID: 19442441.

38. Vila J, Tauler M, Grifoll M. Bacterial PAH degradation in marine and terrestrial habitats. Curr Opin Biotechnol. 2015; 33:95–102. 10.1016/j.copbio.2015.01.006. PubMed PMID: 25658648.

39. Abdel-Shafy HI, Mansour MS. A review on polycyclic aromatic hydrocarbons: source, environmental impact, effect on human health and remediation. Egypt J Pet. 2016; 25:107–23. 10.1016/j.ejpe.2015.03.011.

40. Peng T, Kan J, Hu J, Hu Z. Genes and novel sRNAs involved in PAHs degradation in marine bacteria *Rhodococcus* sp. P14 revealed by the genome and transcriptome analysis. 3 Biotech. 2020; 10:140. 10.1007/s13205-020-2133-6. PubMed PMID: 32206489.

41. Gerdes D. The Pacific oyster *Crassostrea gigas*: Part I. Feeding behaviour of larvae and adults. Aquaculture. 1983; 31:195–219. 10.1016/0044-8486(83)90313-7.

42. Vaezzadeh V, Zakaria MP, Bong CW, Masood N, Mohsen Magam S, Alkhadher S. Mangrove oyster (*Crassostrea belcheri*) as a biomonitor species for bioavailability of polycyclic aromatic hydrocarbons (PAHs) from sediment of the west coast of peninsular Malaysia. Polycyclic Aromat Compd. 2019; 39:470–85. 10.1080/10406638.2017.1348366.

43. Gan N, Martin L, Xu W. Impact of Polycyclic Aromatic Hydrocarbon Accumulation on Oyster Health. Front Physiol. 2021; 12:734463. 10.3389/fphys.2021.734463. PubMed PMID: 34566698; PubMed Central PMCID: PMCPMC8461069.

44. Sarkar A, Bhagat J, Saha Sarker M, Gaitonde DCS, Sarker S. Evaluation of the impact of bioaccumulation of PAH from the marine environment on DNA integrity and oxidative stress in marine rock oyster (*Saccostrea cucullata*) along the Arabian sea coast. Ecotoxicology. 2017; 26:1105–16. 10.1007/s10646-017-1837-9. PubMed PMID: 28755287.

45. Sharma J, Sundar D, Srivastava P. Biosurfactants: Potential Agents for Controlling Cellular Communication, Motility, and Antagonism. Front Mol Biosci. 2021; 8:727070. 10.3389/fmolb.2021.727070. PubMed PMID: 34708073.

46. Guttenplan SB, Kearns DB. Regulation of flagellar motility during biofilm formation. FEMS Microbiol Rev. 2013; 37:849–71. 10.1111/1574-6976.12018. PubMed PMID: 23480406.

47. Prüß BM. Involvement of two-component signaling on bacterial motility and biofilm development. J Bacteriol. 2017; 199:e00259–17. 10.1128/jb.00259-17. PubMed PMID: 28533218.

48. Zan J, Cicirelli EM, Mohamed NM, Sibhatu H, Kroll S, Choi O, et al. A complex LuxR- LuxI type quorum sensing network in a roseobacterial marine sponge symbiont activates flagellar motility and inhibits biofilm formation. Mol Microbiol. 2012; 85:916–33. 10.1111/j.1365-2958.2012.08149.x. PubMed PMID: 22742196.

49. Patriquin GM, Banin E, Gilmour C, Tuchman R, Greenberg EP, Poole K. Influence of quorum sensing and iron on twitching motility and biofilm formation in *Pseudomonas aeruginosa*. J Bacteriol. 2008; 190:662–71. 10.1128/JB.01473-07. PubMed PMID: 17993517.

50. Busi S, Rajkumari J. Biosurfactant: a promising approach toward the remediation of xenobiotics, a way to rejuvenate the marine ecosystem. Mar Pollut Microb Remed. 2017:87–104. 10.1007/978-981-10-1044-6_6.

51. Dias MAM, Nitschke M. Bacterial-derived surfactants: an update on general aspects and forthcoming applications. Braz J Microbiol. 2023; 54:103–23. 10.1007/s42770-023-00905-7. PubMed PMID: 36662441.

52. Spring S, Scheuner C, Goker M, Klenk HP. A taxonomic framework for emerging groups of ecologically important marine gammaproteobacteria based on the reconstruction of evolutionary relationships using genome-scale data. Front Microbiol. 2015; 6:281. 10.3389/fmicb.2015.00281. PubMed PMID: 25914684.

53. Huang KC, Mukhopadhyay R, Wen B, Gitai Z, Wingreen NS. Cell shape and cell-wall organization in Gram-negative bacteria. Proc Natl Acad Sci U S A. 2008; 105:19282–7. 10.1073/pnas.0805309105. PubMed PMID: 19050072.

54. Csonka LN. Physiological and genetic responses of bacteria to osmotic stress. Microbiol Rev. 1989; 53:121–47. 10.1128/mr.53.1.121-147.1989. PubMed PMID: 2651863.

55. Bredeston LM, Marciano D, Albanesi D, De Mendoza D, Delfino JM. Thermal regulation of membrane lipid fluidity by a two-component system in *Bacillus subtilis*. Biochem Mol Biol Educ. 2011; 39:362–6. 10.1002/bmb.20510. PubMed PMID: 21948508.

56. Wen Z, Zhang J-R. Bacterial capsules. Molecular medical microbiology: Elsevier; 2015. p. 33–53. 10.1016/B978-0-12-397169-2.00003-2.

57. Nelson EJ, Harris JB, Morris JG, Jr., Calderwood SB, Camilli A. Cholera transmission: the host, pathogen and bacteriophage dynamic. Nat Rev Microbiol. 2009; 7:693–702. 10.1038/nrmicro2204. PubMed PMID: 19756008.

58. Zhuang GC, Peña-Montenegro TD, Montgomery A, Hunter KS, Joye SB. Microbial metabolism of methanol and methylamine in the Gulf of Mexico: insight into marine carbon and nitrogen cycling. Environ Microbiol. 2018; 20:4543–54. 10.1111/1462-2920.14406. PubMed PMID: 30209867.

59. Jiang X, Jiao N. Nitrate assimilation by marine heterotrophic bacteria. Sci China Earth Sci. 2016; 59:477–83. 10.1007/s11430-015-5212-5.

60. Grein F, Ramos AR, Venceslau SS, Pereira IA. Unifying concepts in anaerobic respiration: insights from dissimilatory sulfur metabolism. Biochim Biophys Acta. 2013; 1827:145–60. 10.1016/j.bbabio.2012.09.001. PubMed PMID: 22982583.

61. An L, Yan YC, Tian HL, Chi CQ, Nie Y, Wu XL. Roles of sulfate-reducing bacteria in sustaining the diversity and stability of marine bacterial community. Front Microbiol. 2023; 14:1218828. 10.3389/fmicb.2023.1218828. PubMed PMID: 37637129.

62. Christensen-Dalsgaard M, Gerdes K. Two higBA loci in the *Vibrio cholerae* superintegron encode mRNA cleaving enzymes and can stabilize plasmids. Mol Microbiol. 2006; 62:397–411. 10.1111/j.1365-2958.2006.05385.x. PubMed PMID: 17020579.

63. Wood TL, Wood TK. The HigB/HigA toxin/antitoxin system of *Pseudomonas aeruginosa* influences the virulence factors pyochelin, pyocyanin, and biofilm formation. Microbiologyopen. 2016; 5:499–511. 10.1002/mbo3.346. PubMed PMID: 26987441.

64. Xue C, Sashital DG. Mechanisms of Type I-E and I-F CRISPR-Cas Systems in *Enterobacteriaceae*. EcoSal Plus. 2019; 8. 10.1128/ecosalplus.ESP-0008-2018. PubMed PMID: 30724156.

65. Mattick JS. Type IV pili and twitching motility. Annu Rev Microbiol. 2002; 56:289–314. 10.1146/annurev.micro.56.012302.160938. PubMed PMID: 12142488.

66. Hospenthal MK, Costa TRD, Waksman G. A comprehensive guide to pilus biogenesis in Gram-negative bacteria. Nat Rev Microbiol. 2017; 15:365–79. 10.1038/nrmicro.2017.40. PubMed PMID: 28496159.

67. Tennent JM, Mattick JS. Type 4 fimbriae. Fimbriae Adhesion, Genetics, Biogenesis, and Vaccines. CRC Press. 2020; p.127-46. 10.1201/9781003068259.

68. Burdman S, Bahar O, Parker JK, De La Fuente L. Involvement of Type IV Pili in Pathogenicity of Plant Pathogenic Bacteria. Genes (Basel). 2011; 2:706–35. 10.3390/genes2040706. PubMed PMID: 24710288.

69. Tekedar HC, Patel F, Blom J, Griffin MJ, Waldbieser GC, Kumru S, et al. Tad pili contribute to the virulence and biofilm formation of virulent *Aeromonas hydrophila*. Front Cell Infect Microbiol. 2024; 14:1425624. 10.3389/fcimb.2024.1425624. PubMed PMID: 39145307.

70. Comolli JC, Hauser AR, Waite L, Whitchurch CB, Mattick JS, Engel JN. *Pseudomonas aeruginosa* gene products PilT and PilU are required for cytotoxicity in vitro and virulence in a mouse model of acute pneumonia. Infect Immun. 1999; 67:3625–30. 10.1128/IAI.67.7.3625-3630.1999. PubMed PMID: 10377148.

71. Persat A, Inclan YF, Engel JN, Stone HA, Gitai Z. Type IV pili mechanochemically regulate virulence factors in *Pseudomonas aeruginosa*. Proc Natl Acad Sci U S A. 2015; 112:7563–8. 10.1073/pnas.1502025112. PubMed PMID: 26041805.

